# Molecular signaling associated with antidepressant actions exhibits diurnal fluctuations in the prefrontal cortex and hippocampus of adult male and female mice

**DOI:** 10.64898/2026.04.07.716906

**Authors:** Gemma González-Hernández, Stanislav Rozov, Esther Berrocoso, Tomi Rantamäki

## Abstract

An increasing number of epidemiological and experimental studies have demonstrated a bidirectional relationship between mood disorders and the circadian system, with disrupted circadian rhythms contributing to depressive states, and their restoration playing a key role in antidepressants effects. In this context, we sought to examine whether key molecular targets of antidepressants exhibit diurnal regulatory patterns. Naïve adult male and female C57BL/6 mice were euthanized at 3-hour intervals beginning at *Zeitgeber Time* 0 (ZT0), and hippocampal (HC) and medial prefrontal cortex (mPFC) tissues were collected for RT-qPCR and western blot analyses. We observed statistically significant diurnal rhythmicity in all analyzed transcripts (*cFos*, *Arc*, *Nr4a1*, *Dusp1*, *Dusp5*, and *Dusp6*) in both HC and mPFC samples, with peak expression occurring during the dark (active) phase (ZT15–18). Phosphorylation levels of TrkB^Y816^ (tropomyosin-related kinase) and GSK3β^S9^ (glycogen synthase kinase 3β) also showed periodic rhythmicity, peaking during the light (inactive) phase. Levels of p-ERK2^T185/Y187^ (extracellular-signal regulated kinase) did not display rhythmicity, but peaked during the light phase in the HC, especially in males. Collectively, these findings demonstrate that antidepressant targets are subject to diurnal regulation, highlighting the importance of integrating circadian biology and time-of-day as relevant variables in the development of translationally relevant antidepressant research.

**Highlights:**

- Key molecular targets of antidepressants exhibit diurnal regulation in adult mice
- Diurnal patterns were conserved across targets, sexes, and brain regions (HC&PFC)
- *cFos, Arc, Nr4a1, Dusp1,5,6* mRNAs display peak expression during the dark phase
- TrkB^Y816^ and GSK3β^S9^ phosphorylation peak during the light (inactive) phase
- Antidepressant mechanisms may be linked with circadian and sleep-wake dynamics

**Graphical abstract:** 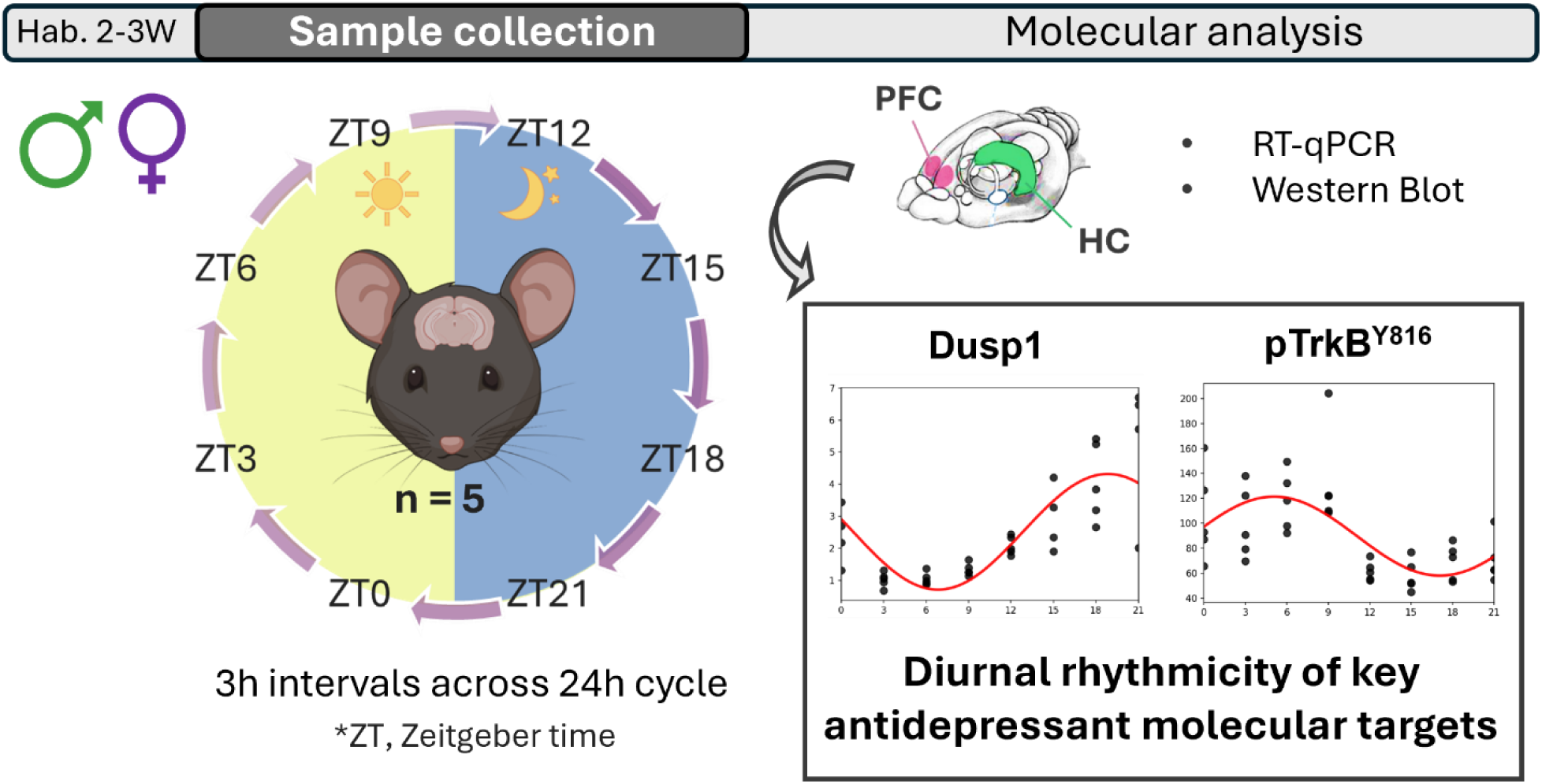

## 1. Introduction

The circadian system controls biochemical and physiological processes that cycle rhythmically over a 24-hour period in alignment with the natural light–dark cycle (de Leeuw et al., 2023). Beyond its canonical role in regulating and maintaining circadian rhythmicity, the circadian system controls a broad range of physiological functions, such as sleep, learning and memory, feeding, and energy balance (de Leeuw et al., 2023; Sato et al., 2022). Notably, an increasing number of epidemiological and experimental studies have demonstrated a bidirectional relationship between the circadian system and mood disorders, with disrupted circadian rhythms contributing to depressive states, and their restoration playing a key role in antidepressants effects (Lyall et al., 2018; Wulff et al., 2010). Indeed, circadian rhythm disruptions have been widely documented in patients and preclinical models of depression, including alterations in sleep-wake patterns, body temperature, hormonal secretion (e.g. cortisol/corticosterone and melatonin), mood, activity and appetite (de Leeuw et al., 2023; Orozco-Solis et al., 2016). Irregular expression patterns of the core genes constituting the molecular circadian clock (i.e. clock genes), as well as of the genes under their direct regulation (i.e. clock-controlled genes), have also been reported (Li et al., 2013).

Rapid-acting antidepressant treatments, most notably subanaesthetic-dose ketamine (aan het Rot et al., 2010; Murrough et al., 2013), can alleviate depressive symptoms within hours after a single or few dose administrations, and their antidepressant effects persist well beyond their pharmacokinetic duration (Witkin et al., 2019). Ketamine, along with other antidepressant therapies, has been shown to modulate neuroplasticity in brain regions implicated in depression, including the medial prefrontal cortex (mPFC) and hippocampus (HC). Activation of BDNF (brain-derived neurotrophic factor) receptor TrkB (tropomyosin-related kinase B) and its downstream pathways, have been proposed as key mechanisms underlying antidepressant-induced neuroplasticity (Casarotto et al., 2021; Castrén & Antila, 2017; Saarelainen et al., 2003). Among downstream pathways, the activation of MAPK (mitogen-activated protein kinase, also known as extracellular-signal regulated kinase; ERK) (Humo et al., 2020; Réus et al., 2014) and inhibition of GSK3β (glycogen synthase kinase 3 β) (Beurel et al., 2016; Li & Jope, 2010), have been especially implicated.

Rapid-acting antidepressants, including pharmacological treatments such as ketamine and non-pharmacological interventions like electroconvulsive therapy and sleep deprivation, engage common molecular pathways. These treatments acutely increase glutamatergic signaling and upregulate immediate-early genes (IEGs; including *Bdnf* and *Arc)* in the mPFC and HC of adult rodents (Kohtala & Rantamäki, 2021; Sato et al., 2022). Nitrous oxide (N_2_O), another putative rapid-acting antidepressant (Nagele et al., 2018), has been shown to similarly increase markers of neuronal excitability in the mPFC, including ERK phosphorylation and the expression of *Dusp1*, *cFos*, *Arc* and *Bdnf* (Kohtala et al., 2019a). Interestingly, psilocybin, a psychedelic compound with rapid antidepressant therapeutic potential, also acutely increased *Dusp1*, *cFos*, and *Nr4a1* expression in the mPFC (Jefsen et al., 2021). Follow-up studies suggest that, following this acute cortical excitation, both pharmacological and non-pharmacological rapid-acting antidepressants modulate TrkB and GSK3β signaling (Alitalo et al., 2023, 2024; Kohtala et al., 2019b). Moreover, a transcriptome-wide study revealed converged regulation of MAPK pathway by both ketamine and N_2_O in the mPFC, primarily affecting the transcription of dual-specificity phosphatases (DUSPs; also known as MAP kinase phosphatases; MKPs) (Rozov et al., 2024), which are important negative regulators of MAPK proteins (Pérez-Sen et al., 2019).

Another transcriptome-wide study also reported that both ketamine and sleep deprivation altered MAPK signaling pathway, including the expression of *Dusp* genes, in the mouse anterior cingulate cortex (Orozco-Solis et al., 2017). The affected pathways in both studies were predominantly associated with synaptic function and neuronal activity, but also biological processes related to circadian regulation of gene expression, immune response, protein synthesis and cellular metabolism - processes that are all governed by the circadian system.

Importantly, GSK3β (Besing et al., 2017; Sahar et al., 2010) and MAPK (Akashi & Nishida, 2000; Goldsmith & Bell-Pedersen, 2013; Wang et al., 2020) are also well-established regulators of the circadian clock, which suggests an overlap or interaction between neuroplasticity and circadian pathways. Recent research highlights the circadian system as a key factor potentially involved in rapid antidepressant responses *via* these shared intracellular signaling cascades, as well as through modulation of sleep and related circadian processes (Kohtala et al., 2021; Lee, 2019; Sato et al., 2022; Wang et al., 2020). Furthermore, we found additional evidence for an interaction between neuroplasticity and circadian mechanisms by performing a preliminary analysis that merged recent RNA-sequencing data of transcripts affected by N_2_O treatment (Rozov et al., 2024), with a genome-wide expression study identifying genes that are diurnally expressed in the mouse PFC (Yang et al., 2007). This analysis yielded a list of transcripts that were both diurnally regulated and modulated by N_2_O, with most significant markers showing synchronized phases **(Supplementary Information File 1).**

In light of growing evidence linking the effects of rapid-acting antidepressants to circadian modulation, and the increasing interest in chronopharmacology as a strategy to enhance antidepressant efficacy (Benedetti et al., 2021; Sato et al., 2022), we explored the hypothesis that key molecular targets of rapid-acting antidepressants are under diurnal regulation. To this end, we assessed the expression and phosphorylation profiles of these targets in the HC and mPFC of naïve male and female C57BL/6 mice, examining whether specific transcripts exhibit diurnal rhythmicity and how protein phosphorylation levels of key signaling pathways fluctuate across the day. By characterizing the temporal dynamics of these molecular targets, this work aims to inform the design of future chronotherapeutic studies and to emphasize time-of-day as an important biological variable for translational antidepressant research.

## 2. Materials and methods

### 2.1. Animals

Eight- to nine-week-old C57BL/6JRccHsd male and female mice (Envigo, Horst, The Netherlands) were used. Experiments in males and females were conducted separately, but all animals were maintained under identical conditions and followed the same experimental protocol. Upon arrival, mice were group-housed (n = 4-5 per cage) under controlled environmental conditions (22 ± 1 °C, 12:12h light−dark cycle, lights on at 6 a.m ∼ *Zeitgeber Time* 0; ZT0) in standard plastic rodent cages containing bedding and nesting material, with *ad libitum* access to food and water. All cages were placed inside a Scantainer (Scanbur, Sweden) located in a dedicated room, isolated from other animal rooms to minimize disturbances from other experimenters or caretakers. Male mice were allowed over two weeks of habituation, whereas females underwent an additional week of habituation to facilitate estrous cycle synchronization (Lee-Boot effect) (Van Der Lee & Boot, 1955). During the habituation period, disturbances were minimized by restricting access to the experimental rooms to the responsible experimenter with whom the mice had been familiarized, and by limiting cage changes to once per week, ensuring that the final cage change occurred at least 4 days before the experimental procedures. At the time of testing, male mice weighed 18–29 g and females 16–20 g. All procedures complied with the European Communities Council Directive 2010/63/EU, and the Society for Neuroscience guidelines, and were approved by the County Administrative Board of Southern Finland (License ESAVI/21911/2022).

### 2.2. Study design and sample collection and processing

After habituation, mice were euthanized at 3-hour intervals throughout a 24-hour cycle (n = 5 per group), generating eight experimental groups, each corresponding to specific circadian time-points: four collected during the light phase (ZT0, ZT3, ZT6 and ZT9) and four during the dark phase (ZT12, ZT15, ZT18 and ZT21). At each designated time-point, one animal at a time was gently removed from the home-cage and transferred to a separate room for rapid cervical dislocation. For dark-phase collections (ZT12-21), these procedures were performed under darkness using night-vision googles to prevent light exposure. Brain extractions and dissections were subsequently carried out in a separated illuminated room. Briefly, bilateral medial prefrontal cortex (mPFC) and hippocampus (HC) were rapidly dissected on a cooled dish and stored at −80 °C until further processing. Left and right regions were collected in separate sample tubes for protein phosphorylation and mRNA analyses.

### 2.3. Quantitative real time PCR (qRT-PCR)

mRNA was extracted using the NucleoSpin RNA Plus Kit (Macherey-Nagel, Germany) according to the manufacturer’s instructions. RNA concentration and purity were assessed with the Nanodrop 2000 Spectrophotometer (Millipore, USA). RNA samples (500 ng) were reverse transcribed into cDNA using the Maxima First Strand cDNA Synthesis Kit (K1672, Thermo Scientific, USA). RT-qPCR was performed on the Lightcycler® 480 System (Roche, Switzerland) using Maxima SYBR Green/ROX qPCR Master Mix (2X) (K0222, Thermo Scientific, USA). The amplification protocol consisted of 45 cycles at 95 °C (15 s), 60 °C (30 s) and 72 °C (30 s). Primer sequences are listed in the **Table S1**. Most custom-made primers had been previously validated and published (Rozov et al., 2024). *Gapdh* and *B-actin* were used as internal reference genes for hippocampal samples, whereas *Rplp0* was used for prefrontal cortex samples. Different reference genes were selected for each brain region because *Gapdh* and *B-actin* showed oscillatory expression across the day in our PFC samples and were therefore unsuitable for normalization. *Rplp0* was chosen based on prior studies identifying it as one of the most stable reference genes in circadian research (Giri & Sundar, 2022; Hadadi et al., 2018; Kosir et al., 2010). Relative mRNA expression (E^-ΔΔCq) of target genes was calculated using the Pfaffl method (Pfaffl, 2001), and expressed relative to ZT3 (ZT3 = 1). PCR amplification efficiency was determined using a serial dilution curve for each transcript and was consistently maintained within the optimal range of 1.8–2.2.

### 2.4. Western Blot

Unilateral HC and mPFC samples were rapidly homogenized and sonicated in a lysis buffer (137 mM NaCl, 20 mM Tris, 1% NP-40, 10% glycerol, 48 mM NaF, H_2_O) containing Pierce^TM^ Phosphatase and Protease Inhibitor Tablets (Thermo Fisher Scientific, Vantaa, Finland). After ∼15 minutes incubation on ice, samples followed two sequential rounds of centrifugation (16000xg, 15 min, +4°C) and the resulting supernatant was collected for further analysis. The total protein concentrations of the samples were measured using DC™ protein assay kit (Bio-Rad Laboratories, Hercules, CA, USA). Samples containing equal amount of total protein (20-25 µg) were mixed in 1:1 ratio with 2x Laemmli Sample Buffer (Bio-Rad) supplemented with β-mercaptoethanol, heated in 100°C for 3 min, and finally loaded to sodium dodecyl sulphate (SDS)–polyacrylamide gels (4–12% NuPAGE™, Thermo Fisher Scientific). Proteins were electrophoretically separated (80V for ∼15min; followed by 150V for ∼80min) under reducing and denaturing conditions using MOPS SDS Running Buffer (NuPAGE™, Invitrogen, Thermo Fisher Scientific). Proteins were then transferred onto a polyvinylidene difluoride (PVDF) membrane using the Trans-Blot Turbo Transfer System (Bio-Rad), following the manufacturer’s instructions. After 1-hour blocking with 5% BSA in Tris-buffered saline with 0.1% Tween (TBST), the membranes were incubated overnight (+4°C) with the following primary monoclonal antibodies: anti-phospho-TrkB^Y816^ (#4168; 1:1000; Cell signaling technology [CST]), anti-phospho-GSK3β^S9^ (#5558, 1:1000, CST), anti-phospho-ERK1/2^T202/Y204^ (#9106, 1:1000 or 1:2000, CST), and anti-Vinculin (#13901, 1:1000, CST). The membranes were washed in TBST and incubated with horseradish peroxidase-conjugated secondary antibodies (1:5000 or 1:10000 in TBST, 1h at RT; Bio-Rad). After subsequent TBST washes, secondary antibodies were visualized using enhanced chemiluminescence (ECL Plus™, Thermo Scientific) for detection with Bio-Rad ChemiDoc™ MP camera (Bio-Rad Laboratories, Helsinki, Finland). Densitometric analysis of the chemiluminescent blots were quantified using ImageLab software (Bio-Rad). Phospho-proteins were normalized against total Vinculin, and results expressed as percentage of ZT3 average, which was set to 100%. Full blots are provided in the **Supplementary Information File 2**.

### 2.5. Sample exclusion criteria

Sample exclusions were applied according to predefined quality criteria for tissue integrity, RNA yield, and assay performance. The initial experimental design included 40 mice per sex (5 per circadian time point), but one male at ZT21 was excluded due to evident brain malformations observed during dissection. Thus, the final number of samples available for downstream analyses was N = 39 for males and N = 40 for females.

Additional exclusions were applied on a dataset-specific basis. Final sample sizes for each dataset are reported in the corresponding figure legends. A detailed description of all sample exclusions and missing values, organized by figure and sex is provided in **Table S2.**

### 2.6. Statistical analysis

RT-qPCR and Western Blot results are presented as relative mRNA expression (EΔΔCq) or protein phosphorylation (% of ZT3), respectively, across the designated circadian time points. Solid lines represent mean values (relative to ZT3, which has been set to 1 or 100% respectively), and light-grey shaded areas indicate the standard deviation (SD). The p-values shown in these graphs correspond to the cosinor rhythmicity analysis performed on our datasets to evaluate the presence of diurnal oscillations.

Cosinor analysis with a fixed 24-hour period was used to model circadian fluctuations by fitting a linearized sinusoidal curve to the data (Cornelissen, 2014). MESOR, amplitude, and acrophase were estimated from the fitted cosine and sine coefficients. In addition, the F statistic, associated p-value and R^2^ were calculated from the F-test to assess model fit and determine statistical significance of the rhythms. A full description of the model is provided in the **Supplementary Methods, Supplementary Information File 2**. Rhythmicity was additionally verified using the Bio_Cycle algorithm (Agostinelli et al., 2016), which yielded comparable results **(Supplementary Information File 3)**. For direct comparisons of peak and through circadian time points in gene expression and protein phosphorylation, either two-tailed unpaired Student’s *t* test or Mann–Whitney *U* tests were applied depending on data normality. Prior to these analyses, outliers were identified using Grubbs’ test (α = 0.01), and data distribution was assessed with Shapiro-Wilk normality test. Data are presented as Mean Value ± SD. Most statistical analyses and graphing were performed using GraphPad Prism v.10 (GraphPad Software, San Diego, CA, USA). A p-value ≤ 0.05 was considered statistically significant in all tests (*p ≤ 0.05, **p ≤ 0.01, ***p ≤ 0.001, ****p ≤ 0.0001). Detailed information on the statistical tests applied in each case is provided in the corresponding figure legends and **Tables S3-6**.

## 3. Results

### 3.1. *c-Fos, Arc, Nr4a1 Dusp1, Dusp5* and *Dusp6* are diurnally regulated in the mouse hippocampus and medial prefrontal cortex

Given the more direct multisynaptic neural connections between the suprachiasmatic nucleus (SCN) and the hippocampus (HC) compared to the medial prefrontal cortex (mPFC) (Ma & Morrison, 2025; Ruby, 2021), we first focused our analyses on the HC to investigate whether transcripts associated with antidepressant effects exhibit fluctuations throughout the day across the 24-hour cycle. All selected markers (*c-Fos, Arc, Nr4a1, Dusp1, Dusp5*, *Dusp6,*) presented diurnal fluctuations and followed a similar pattern of expression, with trough expression during the light phase (ZT3-6) and peak expression during the dark phase (ZT15-18) **(Figure 1).** Indeed, all transcripts demonstrated significant 24-hour rhythmicity, as indicated by the cosinor analysis **(Figure S1).** This was observed in both males **(Figures 1A, S1A)**: ***cFos*** p = 0.043, ***Arc*** p = 6.07×10⁻^5^, ***Nr4a1*** p = 0.0012, ***Dusp1*** p = 0.0149, ***Dusp5*** p = 0.0025, ***Dusp6*** p = 2.52×10⁻^7^; and females **(Figures 1B, S1B): *cFos*** p = 6.9×10⁻^5^, ***Arc*** p = 6.95×10⁻^5^, ***Nr4a1*** p = 0.002, ***Dusp1*** p = 9.68×10⁻^5^, ***Dusp5*** p = 0.0158, ***Dusp6*** p = 2.04×10⁻^6^. Full cosinor output parameters are provided in **Supplementary Information File 4**.

**Figure 1.**
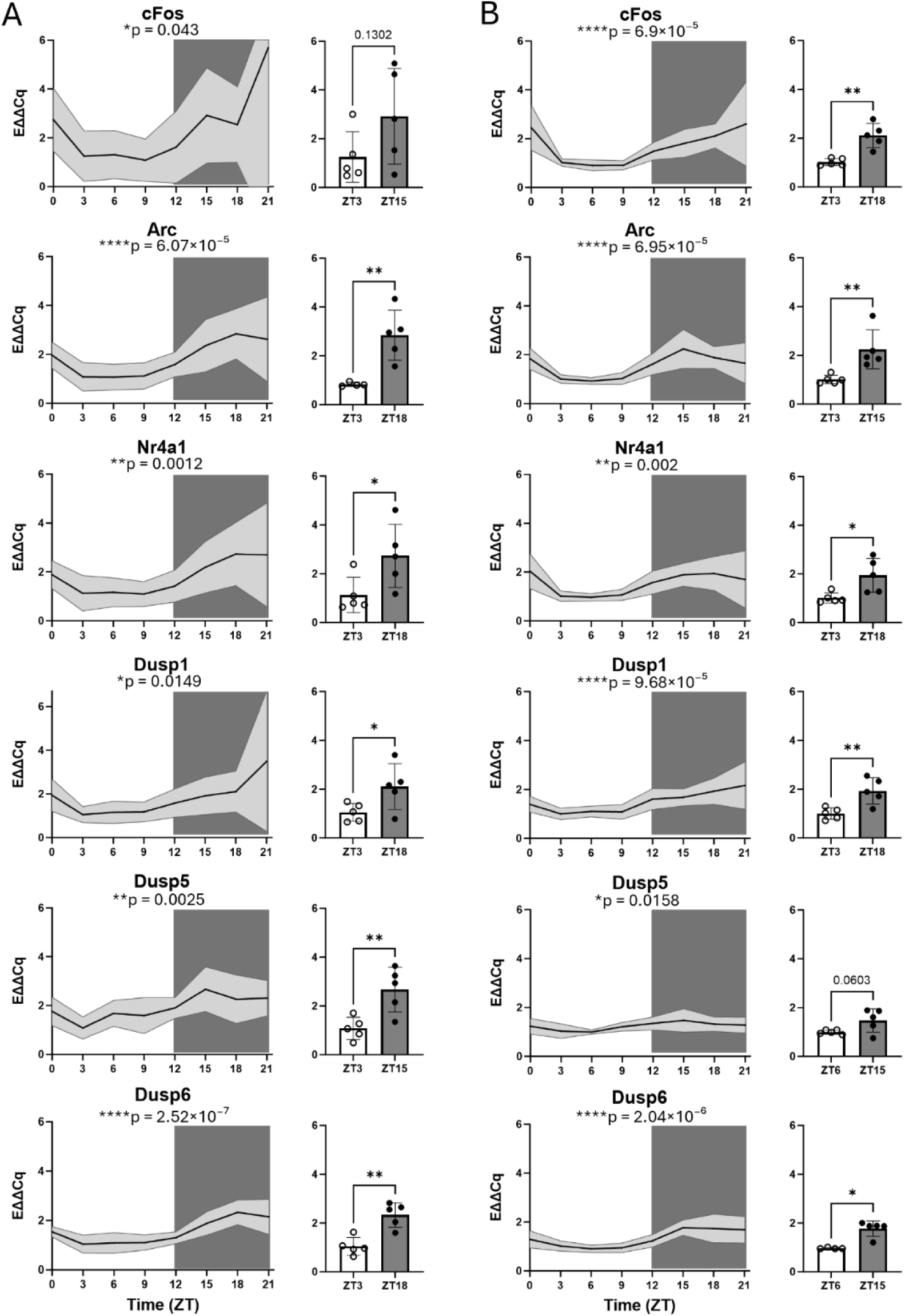
Diurnal regulation of *c-Fos, Arc, Nr4a1, Dusp1, Dusp5*, *Dusp6,* in the male (A) and female (B) mouse hippocampus. Graphs depict relative gene expression levels across an entire 24-hour cycle in 3-hours intervals, alongside direct comparisons between peak and trough time points (ZT3-6 vs. ZT15-18). In the main graphs, solid lines represent mean values (relative to ZT3 = 1), and light-grey shaded areas indicate standard deviation (SD). The p-values indicate the statistical significance of the detected rhythms as determined by the cosinor analysis. Gene expression was quantified by RT-qPCR. Y-axis represents relative expression levels (EAACq) normalized to the geometric mean of *Gapdh* and *Bactin*. X-axis indicates circadian time-points (ZT) of sample collection. For the peak-trough comparisons, Student’s two-tailed unpaired *t* test was used for all targets, except male *Nr4a1* and female *Dusp6*, for which the non-parametric Mann-Whitney test was applied. n = 4-5 per time-point. Data are presented as mean ± SD. * ≤ 0.05, ** ≤ 0.01, *** ≤ 0.001, **** ≤ 0.0001. Detailed statistical information and sample sizes are provided in **Table S3**.

In the mPFC, the cosinor analysis likewise revealed significant rhythmicity for all markers **(Figures 2, S2).** This was again observed in both males **(Figures 2A, S2A): *cFos*** p = 3.2×10⁻^5^, ***Arc*** p = 4.05×10⁻^7^, ***Nr4a1*** p = 5.16×10⁻^6^, ***Dusp1*** p = 2.82×10⁻^5^, ***Dusp5*** p = 5.37×10⁻^8^, ***Dusp6*** p = 1.71×10⁻^10^; and females **(Figure 2B, S2B): *cFos*** p = 5.14×10⁻^8^, ***Arc*** p = 3.78×10⁻^8^, ***Nr4a1*** p = 8.70×10⁻^7^, ***Dusp1*** p = 6.45×10⁻^8^, ***Dusp5*** p = 1.87×10⁻^7^, ***Dusp6*** p = 2.72×10⁻^9^. Full cosinor output parameters are provided in **Supplementary Information File 4**. Notably, the fluctuations in the mPFC appeared highly pronounced, and the diurnal expression profiles of most markers mirrored those observed in the HC, with the lowest expression levels during the light-phase (ZT3) and the highest levels during the dark period (ZT18). Interestingly, in males, expression levels began to decline as early as ZT21 after peaking at ZT18, whereas in females, most markers appeared to still be reaching a plateau around ZT21 and remained elevated until ZT0, followed by a rapid decline upon lights onset.

**Figure 2.**
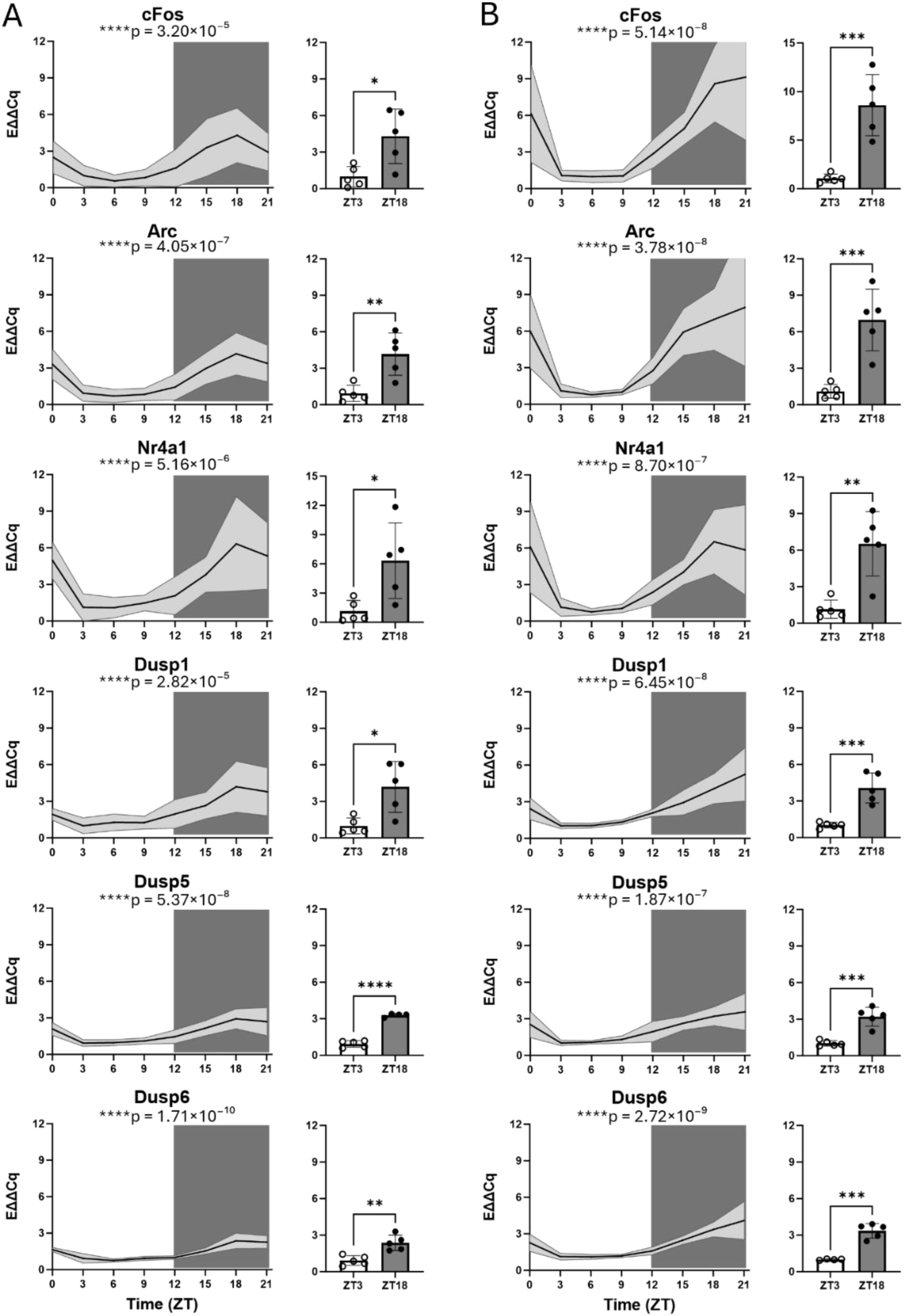
Diurnal regulation of *c-Fos, Arc, Nr4a1, Dusp1, Dusp5*, *Dusp6,* in the male (A) and female (B) mouse prefrontal cortex. Graphs depict relative gene expression levels across an entire 24-hour cycle in 3-hours intervals, alongside direct comparisons between peak and trough time points (ZT3 vs ZT18). In the main graphs, solid lines represent mean values (relative to ZT3 = 1), and light-grey shaded areas indicate standard deviation (SD). p-values correspond to the statistical significance of the detected rhythms as determined by cosinor analysis. Gene expression was quantified by RT-qPCR. Y-axis represents relative expression levels (EAACq) normalized to *Rplp0*. X-axis indicates circadian time-points (ZT) of sample collection. For the peak-trough comparisons, Student’s two-tailed unpaired *t* test was applied. n = 4-5 per time-point. Data are presented as mean ± SD. * ≤ 0.05, ** ≤ 0.01, *** ≤ 0.001, **** ≤ 0.0001. Detailed statistical information and sample sizes are provided in **Table S4**.

In summary, the selected molecular targets of antidepressants displayed prominent diurnal rhythmicity in gene expression, with highly consistent patterns across markers, brain regions and sexes: trough expression during the light phase (ZT3-6) and peak expression during the dark phase (ZT15-18).

### 3.2. TrkB signaling exhibits diurnal rhythmicity and peaks during the inactive period of mice

Given the relevant role of TrkB signaling in depression and antidepressant research (Casarotto et al., 2021; Castrén & Antila, 2017; Kohtala & Rantamäki, 2021; Saarelainen et al., 2003), we next examined whether phosphorylation of TrkB at the Y816 site (pTrkB^Y816^) also undergoes diurnal regulation in the HC **(Figure 3)** and the mPFC **(Figure 4).** This site was selected because it has been demonstrated that pharmacologically diverse antidepressants rapidly and selectively induce phosphorylation at TrkB^Y816^, but not at the Shc binding site (TrkB^Y515^) (Rantamäki et al., 2007).

**Figure 3.**
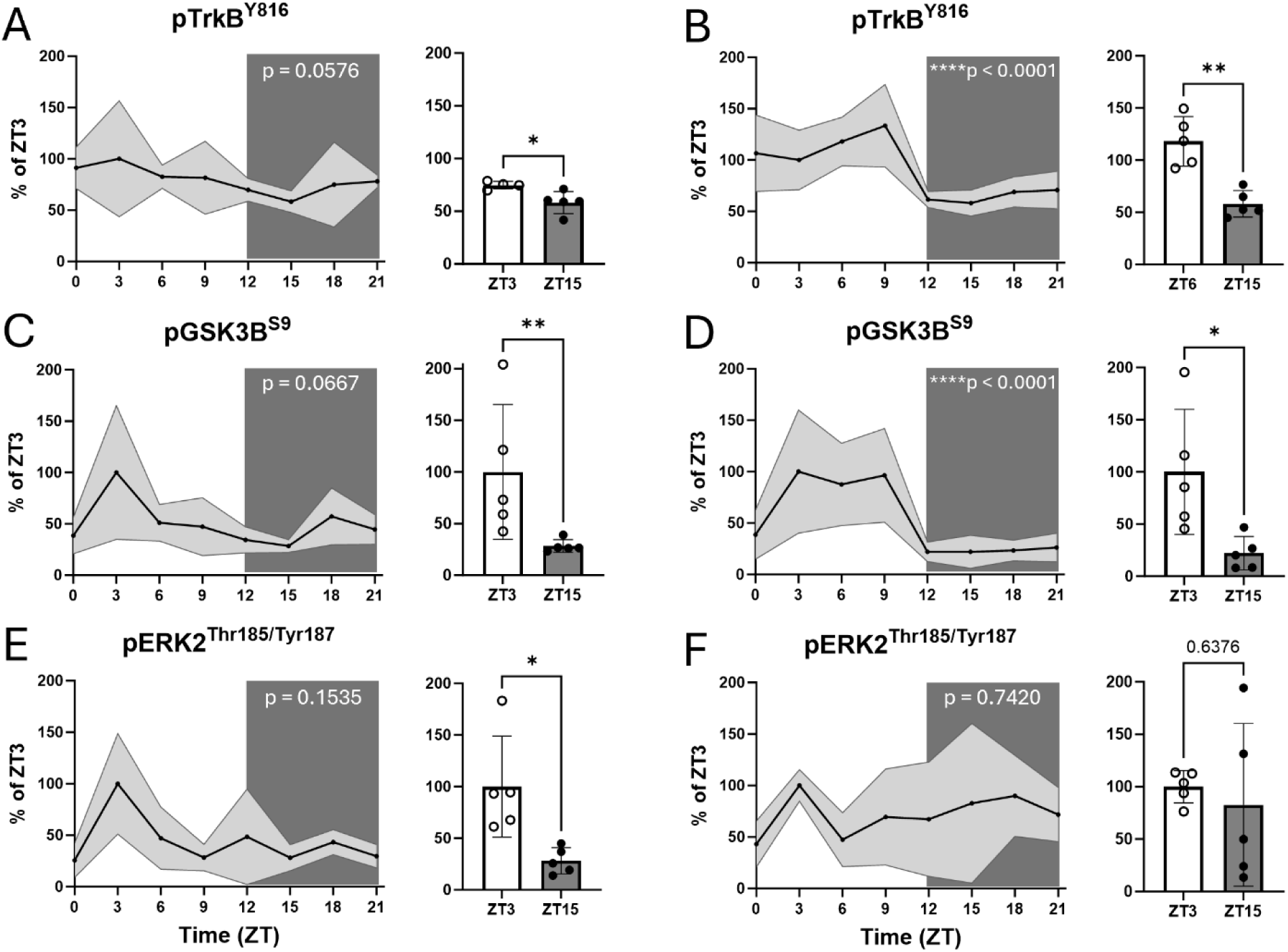
TrkB, GSK3β and ERK2 phosphorylation patterns throughout the 24-hour cycle in the hippocampus of male (A, C, E) and female mice (B, D, **F)**. Left column represents males and right column females. Graphs show the immunoblotting representation of **(A-B)** pTrkB, **(C-D)** pGSK3β, **(E-F)** pERK2 across an entire 24-hour cycle in 3-hours intervals, alongside direct comparisons between peak and trough time points (ZT3-6 vs. ZT15). In the main graphs, solid lines represent mean values (relative to ZT3 = 100%), and light-grey shaded areas indicate standard deviation (SD). The p-values indicate the statistical significance of the detected rhythms as determined by the cosinor analysis. Phosphorylation levels were quantified by western blotting. Y-axis represents phosphoproteins levels normalized against total Vinculin and compared to the average signal of ZT3, set to 100% (expressed as % of ZT3). X-axis indicates circadian time-points (ZT) at which the samples were collected. For the peak-trough comparisons, Student’s two-tailed unpaired *t* test was used for all targets, except male pGSK3B, for which the non-parametric Mann-Whitney test was applied. n = 4-5 per time-point. Data are presented as mean ± SD. * ≤ 0.05, ** ≤ 0.01, *** ≤ 0.001, **** ≤ 0.0001. Detailed statistical information and sample sizes are provided in **Table S5**.

**Figure 4.**
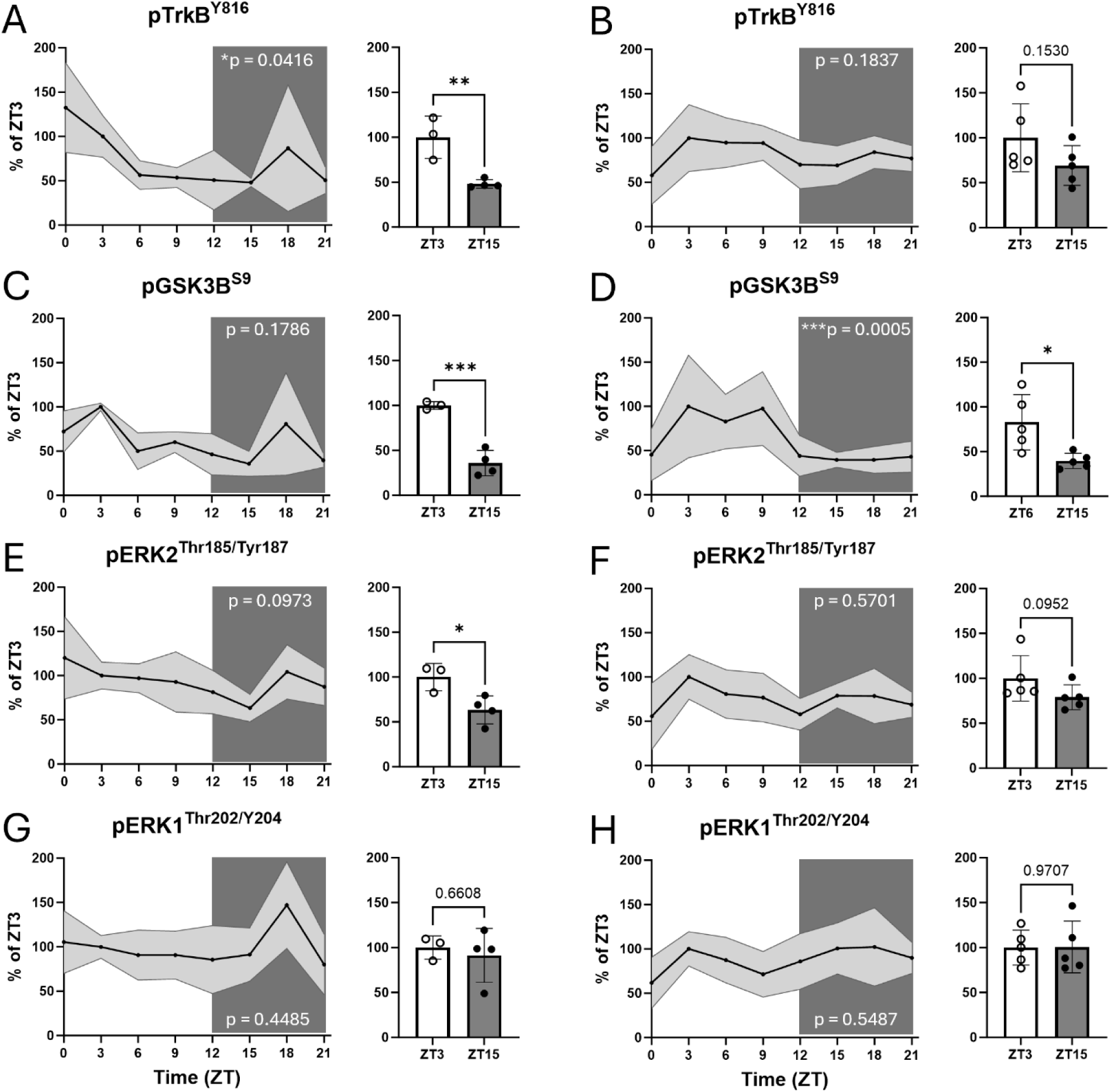
TrkB, GSK3β and ERK1/2 phosphorylation patterns throughout the 24-hour cycle in the prefrontal cortex of male (A, C, E, G) and female mice (B, D, F, H). Left column represents males and right column females. Graphs show the immunoblotting representation of **(A-B)** pTrkB, **(C-D)** pGSK3β, **(E-F)** pERK2, **(G-H)** pERK1 across an entire 24-hour cycle in 3-hours intervals, alongside direct comparisons between peak and trough time points (ZT3-6 vs. ZT15). In the main graphs, solid lines represent mean values (relative to ZT3 = 100%), and light-grey shaded areas indicate standard deviation (SD). The p-values indicate the statistical significance of the detected rhythms as determined by the cosinor analysis. Phosphorylation levels were quantified by western blotting. Y-axis represents phosphoproteins levels normalized against total Vinculin and compared to the average signal of ZT3, set to 100% (expressed as % of ZT3). X-axis indicates circadian time-points (ZT) at which the samples were collected. For the peak-trough comparisons, Student’s two-tailed unpaired *t* test was used for all markers, except female pERK2, for which the non-parametric Mann-Whitney test was applied. n = 3-5 per time-point. Data are presented as mean ± SD. * ≤ 0.05, ** ≤ 0.01, *** ≤ 0.001, **** ≤ 0.0001. Detailed statistical information and sample sizes are provided in **Table S6**.

Overall, pTrkB^Y816^ exhibited clear fluctuations across the day in both brain regions, with significant differences between peak and trough phosphorylation levels, and higher pTrkB levels occurring during the light phase. Cosinor analysis revealed mild rhythmicity in males, with a trend towards statistical significance in the HC (p = 0.0576; **Figure 3A, S3A)** and significant rhythmicity in the mPFC (p = 0.0416; **Figure 4A, S4A).** In contrast, robust and highly significant rhythmicity was observed in the HC of female mice (p < 0.0001; **Figure 3B, S3B).** Although the female mPFC displayed a similar diurnal pattern of TrkB phosphorylation (i.e. higher pTrkB^Y816^ levels during the light phase), the oscillation amplitude was insufficient to reach statistical significance (p = 0.1837; **Figure 4B, S4B**), consistent with the non-significant comparison between ZT3 and ZT15 time points. Despite the lack of statistical significance, the observed trend suggests a similar underlying rhythmicity. Full cosinor output parameters are provided in **Supplementary Information File 4**.

### 3.3. GSK3β and ERK phosphorylation patterns throughout the 24-hour cycle

We then explored selected key downstream signaling pathways linked to the antidepressant actions of ketamine and other rapid-acting treatments, as well as to the regulation of the molecular circadian clock. First, we measured phosphorylation of GSK3β at Ser9 (pGSK3β^S9^), an inhibitory site whose phosphorylation suppresses kinase activity and therefore serve as an inverse indicator of GSK3β function (i.e. higher pGSK3β^S9^ = lower kinase activity). Similarly to phosphorylated TrkB, pGSK3β^S9^ levels also fluctuated across the 24-hour cycle, exhibiting clear and significant differences between peak and trough values, with higher phosphorylation levels during the light phase **(Figures 3, 4)**. In males, cosinor analysis indicated a trend toward significance in the HC (p = 0.0667; **Figures 3C, S3C**), but failed to reach significance in the mPFC (p = 0.1786; **Figures 4C, S4C**). The observed diurnal pattern and rhythmicity were more pronounced in females, displaying robust significance in both the HC (p < 0.0001; **Figures 3D, S3D**) and the mPFC (p = 0.0005; **Figures 4D, S4D**). Full cosinor output parameters are provided in **Supplementary Information File 4**.

Next, we assessed phosphorylation of ERK1/2 at their conserved activation loops residues, Thr202/Tyr204 (ERK1) and Thr185/Tyr187 (ERK2), sites required for full kinase activation and subsequent downstream signaling. Phosphorylated ERK1/2 did not display diurnal rhythmicity based on the cosinor analysis (all p-values > 0.05) **(Figures S3, S4)**. Cosinor output parameters are provided in **Supplementary Information File** In the HC, pERK2^T185/Y187^ appeared to peak at ZT3 **(Figure 3E-F)**, whereas in the mPFC it tended to show increased phosphorylation during both the light and dark phases, suggesting a biphasic profile of phosphorylation, especially in females **(Figure 4E-F).** pERK1^Thr202/Y204^ showed a similar pattern to pERK2 in the mPFC **(Figure 4G-H),** but it was undetectable in the hippocampal samples.

In summary, phosphorylation levels fluctuated throughout the day in both the HC and mPFC, although these diurnal patterns were generally less pronounced than those observed at the transcriptional level. Overall pTrkB and pGSK3β levels were rhythmic and upregulated during the mice inactive period (light phase) in both brain regions. In contrast, pERK2 did not show a clear distinction between light and dark phases in the mPFC, whereas in the HC it appeared to peak around ZT3, more prominently in males. Taken together, diurnal phosphorylation profiles were comparable across brain regions and, to some extent, between sexes; however, females exhibited more pronounced fluctuations and stronger evidence of rhythmicity.

## 4. Discussion

While it is well-known that physiological functions such as hormonal secretion, body temperature or mood fluctuate throughout the day, it is often overlooked in antidepressant research that molecular processes are also governed by circadian (i.e. internal 24-h clock-sustained cycle) and diurnal rhythms (i.e. daily pattern tied to day-night cycle) (Alitalo et al., 2021; Nelson et al., 2021). These fluctuations can affect experimental targets of interest and may act as confounding factors in data interpretation. Here we demonstrate that several molecular targets intimately associated with antidepressant responses exhibit diurnal regulation in the hippocampus (HC) and medial prefrontal cortex (mPFC) of adult naïve mice. Remarkably, the oscillation patterns were largely consistent across markers, sexes, and brain regions, suggesting a fundamental, conserved rhythmic regulation of antidepressant target molecules.

All selected transcripts (*cFos*, *Arc*, *Nr4a1, Dusp1, Dusp5* and *Dusp6*) exhibited significant 24-hour rhythmicity. Notably, a previous genome-wide study reported that ∼10% of transcripts in the mouse PFC exhibit diurnal variation – although, among our targets, only *Dusp6* was identified (Yang et al., 2007). However, our findings align well with more recent RNA-sequencing data showing that these targets are part of the ∼12% of rhythmic transcripts in the mouse mPFC, with *Arc* and *Nr4a1* ranking among the top 10 rhythmic genes based on statistical significance (Burns et al., 2025a). Moreover, *Homer1a*, another target associated with rapid antidepressant-induced plasticity and circadian regulation, has also been shown to follow a similar diurnal expression pattern (Sarrazin et al., 2024), and was likewise identified among the rhythmic transcripts by Burns *et al* (2025a). Expression levels of *Dusps*, and, as expected, expression levels of IEGs (*cFos*, *Arc* and *Nr4a1*) peaked during the dark phase, which corresponds to the active period of mice. Overall, males and females displayed similar expression patterns, however, in male PFC, most targets appeared to peak at ZT18 and then gradually decline, whereas in females, elevated expression levels were sustained from ZT18 to ZT21 and remained high at ZT0, followed by a rapid and pronounced decrease after light onset. These differences likely reflect sex-specific activity profiles, as female mice generally exhibit higher and more sustained levels of exploratory and ambulatory activity, especially during the dark phase (Khatiz et al., 2025; Lightfoot, 2008). In any case, the sharp shifts in expression following lights-on (decrease) and lights-off (increase) indicate that these genes are highly responsive to light cues.

Overall, the phosphorylation levels of the analyzed proteins also fluctuated across the day in both the HC and PFC, as indicated by the significant differences between their trough and peak levels. However, the diurnal patterns and rhythmicity were not as pronounced or consistent as those observed in the gene expression data. TrkB^Y816^ phosphorylation, which is very commonly associated with antidepressant drug responses (Casarotto et al., 2021; Castrén & Antila, 2017; Castrén & Monteggia, 2021; Rantamäki et al., 2007; Saarelainen et al., 2003), showed significant (or a trend towards) rhythmicity in the HC and mPFC of both male and female mice, with higher levels during the light phase, which corresponds to the inactive period of mice. This aligns with previous research linking antidepressant-induced TrkB signaling to homeostatic emergence of deep sleep and associated physiological alterations (e.g. drop in temperature) (Alitalo et al., 2023; ElGrawani et al., 2024). Although we did not address molecular mechanisms underlying observed changes in TrkB phosphorylation, earlier evidence indicates that BDNF displays circadian rhythmicity under constant darkness conditions in the suprachiasmatic nucleus (SCN) and other brain regions, with BDNF protein typically peaking during the subjective night and mRNA peaking in the early subjective day (Liang et al., 1998; Liang et al., 2025). Moreover, Coria-Lucero et al., (2016) reported that *Bdnf* and *TrkB* transcripts in the PFC show circadian regulation in constant darkness, with *Bdnf* exhibiting an ultradian pattern (peaks in both subjective day and night) and *TrkB* peaking during the subjective night (active phase). The authors proposed that this regulation may involve direct BMAL1:CLOCK control, supported by the presence of E-box and CRE sites in their promoters.

GSK3β is an important modulator of the molecular clock. It directly phosphorylates core clock components and regulates BMAL1 protein stability, thereby influencing the amplitude and timing of circadian oscillations (Sahar et al., 2010). Moreover, GSK3β phosphorylation has been shown to follow a circadian rhythm with peaks during the subjective day -inactive phase- in male SCN, influencing neuronal excitability and plasticity, as well as regulating clock gene periodicity (Besing et al., 2015; Paul et al., 2016). Besing et al. (2017) further demonstrated 24-hour oscillations in hippocampal pGSK3β levels under standard light–dark conditions, with higher phosphorylation during the light phase. Importantly, this rhythm persisted under constant darkness, confirming that it is driven by the endogenous circadian clock even in the absence of external light cues. Consistent with these findings, our pGSK3β data reveal a comparable diurnal phosphorylation profile, with higher levels during the light phase, as also seen with pTrkB. Importantly, we extend previous observations by demonstrating that this diurnal regulation also occurs in the PFC, and by showing that the oscillatory pattern is present in both sexes but is particularly pronounced in females. The rhythmic phosphorylation patterns observed for TrkB and GSK3β may reflect time-of-day–dependent modulation of kinase activity, with implications for both neuronal plasticity and circadian regulation.

ERK1/2 modulates synaptic plasticity, mood, learning and memory (Eckel-Mahan et al., 2008; Thomas & Huganir, 2004), and it’s a key modulator of the molecular circadian clock (Akashi & Nishida, 2000; Goldsmith & Bell-Pedersen, 2013; Wang et al., 2020). Preclinical and clinical evidence implicates ERK signaling in depression (Dwivedi et al., 2001; Yuan et al., 2010), and rapid antidepressant responses (Pochwat et al., 2017; Qi et al., 2009; Réus et al., 2014; Wang et al., 2020). Phosphorylated ERK has been shown to influence circadian gene expression through direct interactions with core clock proteins (e.g. CLOCK, BMAL1, CRY1/2) and *via* downstream effectors and transcription factors such as CREB (Goldsmith & Bell-Pedersen, 2013; Oh-hashi et al., 2002; Roux & Blenis, 2004; Sanada et al., 2002; Travnickova-Bendova et al., 2002). Notably, ERK1/2 phosphorylation has been reported to exhibit circadian rhythmicity in the hamster SCN, peaking in the (subjective) day under both light-dark and constant conditions, and showing photic induction at night (Coogan & Piggins, 2003). Furthermore, the MAPK/ERK pathway has been identified as a key mechanism by which light exposure induces IEG expression, such as *cFos*, in the SCN, translating photic stimuli into molecular responses (Dziema et al., 2003). In the hippocampus, pERK1/2 also displays circadian oscillations in male mice under both light-dark cycles and constant darkness, with peak levels occurring during the light phase (∼ZT4) (Eckel-Mahan et al., 2008). Collectively, this evidence supports a role for ERK signaling in circadian regulation and in the resetting of circadian rhythms by both photic and non-photic stimuli (Coogan & Piggins, 2003; Wang et al., 2020). Consistent with these reports, our hippocampal pERK2 data tended to be higher during the light phase in males, although no statistically significant rhythmicity was detected. Females also showed a peak at ZT3, however, greater variability during the dark phase obscured a clear light-phase increase. In the mPFC, pERK2 patterns were less consistent, with light-phase elevation in males and a biphasic profile in females. Whether these regional and sex-dependent differences reflect true biological variation remains uncertain.

The intensity and duration of MAPK signaling, including ERK1/2, is tightly regulated by DUSP phosphatases. *Dusp* genes have also been associated with several mental and neurological disorders (An et al., 2021; Pérez-Sen et al., 2019). Whole-genome expression profiling of postmortem brain tissue, has shown that DUSP1, also known as MKP-1, a nuclear phosphatase that dephosphorylates and inactivates ERK1/2, among others, is overexpressed in subjects with major depressive disorder (Duric et al., 2010). Preclinical studies further reveal that stress- and pain-induced depression in mice upregulates *Dusp1* and downregulates p-ERK levels in the anterior cingulate cortex and HC (Barthas et al., 2017; Humo et al., 2020; Vlasov et al., 2023). Additionally, the *Dusp1* gene has been associated with depression, anxiety and pain (Descalzi et al., 2017). Evidence for circadian or diurnal regulation of *Dusps* was limited to whole-transcriptome analyses showing that some *Dusp* genes are expressed in a circadian manner mainly across peripheral tissues in mice and primates (Mure et al., 2018; Zhang et al., 2014). Of note, *Dusp1* has been also previously identified as the first phosphatase gene shown to function both in a light-dependent and a time-of-day–dependent manner within the mammalian central clock (Doi et al., 2007). Our findings provide direct evidence that *Dusp* expression is indeed diurnally regulated in the HC and PFC, in a consistent manner across all targets tested (*Dusp1*, *Dusp5* and *Dusp6*).

The observed diurnal rhythmicity of rapid-acting antidepressant molecular targets has important implications for preclinical research and translational drug discovery. Although many antidepressant therapies modulate these signaling pathways, findings are often inconsistent across studies, likely due to unreported differences in dosing, sampling time, and the circadian time (ZT) at which treatments are administered. Failure to account for circadian biology, including the opposing circadian nature of nocturnal rodents and diurnal humans, compromises reproducibility and limits translational relevance, particularly for oscillatory targets that may not be conserved across species. Identifying diurnal rhythmicity also opens opportunities for chronotherapeutic optimization, as antidepressant efficacy may depend on the circadian phase at which treatment is delivered due to time-dependent fluctuations in target signaling. Ignoring this temporal dimension may contribute to interindividual variability in treatment response and lead to underestimated efficacy, particularly in psychiatric disorders such as depression, which are characterized by disrupted circadian rhythms. The present work identified trough and peak expression time-points for key molecular targets, which can be used to guide future preclinical chronopharmacological experiments. Given the expected relevance of these targets for antidepressant effects, dosing at peak expression may potentially maximize therapeutic outcomes compared to trough administration. From a translational perspective, this peak time point may also be more optimal, as it coincides with the active phase of rodents, which more closely corresponds to the typical timing of treatment administration in patients. Understanding how molecular targets oscillate throughout the day and how these rhythms are altered in pathophysiological conditions may inform the development of combined pharmacological and circadian interventions (e.g. light therapy, sleep manipulation, timed dosing) to enhance and prolong antidepressant efficacy. These considerations highlight the need to treat time-of-day as a biologically meaningful mechanistic variable in preclinical and early-phase research and to integrate chronopharmacology into drug development pipelines in order to improve efficacy, reduce side effects and support personalized antidepressant treatments.

The inclusion of both sexes in our study is a major strength, given that most preclinical biomedical research, including most of the background literature of this study, focuses exclusively on males despite well-documented sex differences in circadian rhythmicity, gene expression and molecular signaling. Females generally show greater circadian flexibility and sensitivity to light, while males and females differ in the timing and regional distribution of rhythmic transcripts in the brain (Burns et al., 2025a; Lordan, 2025; Vidafar et al., 2024; Walton et al., 2022). Sex-specific modulation of signaling pathways, such as DUSP6–ERK, further suggests that male and female brains may respond differently to the same molecular perturbations. In particular, DUSP6 interacts with sex-specific hormonal and signaling environments to differentially regulate neuroplasticity, stress responses, and cognition, with some studies suggesting a sex-dependent role of DUSP6 in depressive disorder and other psychiatric conditions (An et al., 2021). Accordingly, although we observed relatively similar gene expression levels between sexes in naïve mice, sex differences might become apparent only under specific interventions, stressors or disease states. Although the mechanisms underlying sex differences in transcriptomic rhythms remain unclear, these differences may be driven by sex hormone–dependent modulation of circadian phase and amplitude in clock genes and their downstream targets (Burns et al., 2025a). Sex differences in neuronal cortical activity, including lower daytime firing rates and reduced spike frequency adaptation and action potential amplitude in females during the dark phase, may further contribute to distinct gene expression dynamics (Burns et al., 2025b; Kuljis et al., 2013). Given growing evidence supporting the role of gonadal hormones and the estrous cycle in circadian plasticity and phase-shifting, it is important to mention that estrous stage was controlled in this study *via* vaginal cytology; however, female data were pooled, as most animals were synchronized in the estrus/proestrus stage.

Diurnal expression patterns of the selected targets were largely conserved across brain regions and sexes. Notably, we did observe region- and sex-specific differences in reference gene stability across the 24-hour cycle. Specifically, *Gapdh* and *B-actin* showed stable expression in the HC but not in the mPFC, while *Rplp0* was highly stable in the female mPFC, but showed a gradual increase throughout the day in males. To address potential normalization bias, we performed an alternative analysis where mRNA levels of target transcripts were expressed relative to a common calibrator (male and female ZT3 average = 1) and normalized to the geometric mean of male and female *Rplp0* values at each circadian time-point (**Figure S5**).

Future research should investigate the contribution of these targets to antidepressant responses, examining their diurnal and circadian regulation at transcriptional, protein, and posttranslational levels. Studies should be extended to animal models of depression and incorporate region- and cell-type–specific analyses, as rhythmicity can differ across brain regions and cell populations. Given that these oscillatory patterns vary with sex and age (Walton et al., 2022), further work is needed to elucidate mechanistic sex- and age-dependent differences to inform the development of tailored therapeutic strategies.

## 5. Conclusions

This work demonstrates that molecular targets intimately connected with antidepressant responses are subject to diurnal regulation, with consistent oscillatory patterns across markers, sexes and brain regions indicating a fundamental, conserved rhythmic regulatory mechanism. Sharp shifts in expression following lights onset / offset further suggest a strong diurnal component. These dynamic patterns support the need for personalized (e.g. sex-specific), time-informed treatment strategies, and underscore chronopharmacology as a critical consideration for clinical applications. Overall, our findings emphasize the importance of integrating circadian knowledge into neuroscience research and considering diurnal rhythmicity when investigating antidepressant mechanisms and developing novel psychiatric treatments.

## Author Contributions

G.G-H., S.R and T.R. planned the experiments; G.G-H. and S.R. carried out research; G.G-H. prepared the figures; G.G-H. and S.R. run statistical tests, G.G-H and T.R. wrote the original version of the manuscript; G.G., T.R. and E.B. provided funding; all authors commented the manuscript and accepted final submitted version.

## Ethical statement

All animal experiments were carried out according to the National Institute of Health (NIH) guidelines for the care and use of laboratory animals and approved by National Animal Experiment Board of (License number ESAVI/21911/2022).

## Acknowledgements

We would like to thank Liisa Konttinen and personnel of the Laboratory Animal Centre of the University of Helsinki (Biocenter 3) for their technical assistance. We thank Dr. Marion Schott, Dr. Ipek Yalcin, and Dr. Michel Barrot from the University of Strasbourg for sharing the custom-designed Dusp1 primers. We are grateful to Gerard Sánchez Maltas for his valuable assistance and technical support with the implementation of cosinor analysis. Dr. Samuel Kohtala, Iina Annala and other members of the KohtaLab and Laboratory of Neurotherapeutics are thanked for all valuable comments and discussions.

