## Supplementary Information File 2 for "Molecular signaling associated with antidepressant actions exhibits diurnal fluctuations in the prefrontal cortex and hippocampus of adult male and female mice"

**List of Supplementary Material:**

##### Supplementary Methods:

Cosinor Analysis – Formulation and Rationale

##### Supplementary Figures

**Figure S1.** Diurnal rhythmicity of *c-Fos, Arc, Nr4a1, Dusp1, Dusp5*, *Dusp6,* in the male (A) and female (B) mouse hippocampus.

**Figure S2.** Diurnal rhythmicity of *c-Fos, Arc, Nr4a1, Dusp1, Dusp5*, *Dusp6,* in the male (A) and female (B) mouse prefrontal cortex.

**Figure S3.** TrkB, GSK3β and ERK2 phosphorylation patterns throughout the 24-hour cycle in the hippocampus of male (A, C, E) and female mice (B, D, F).

**Figure S4.** TrkB, GSK3β and ERK1/2 phosphorylation patterns throughout the 24-hour cycle in the prefrontal cortex of male (A, C, E, G) and female mice (B, D, F, H).

**Figure S5.** Diurnal regulation of *c-Fos, Arc, Nr4a1, Dusp1, Dusp5, Dusp6*, in the male and female mouse prefrontal cortex – Additional analysis.

##### Supplementary Tables

**Table S1.** Primers used for RT-qPCR.

**Table S2.** Description of sample exclusions and missing values by figure and sex.

**Table S3-6.** Summary of statistical analyses applied in Figures 1-4.

##### Full Blots

### Supplementary Methods

#### Cosinor Analysis – Formulation and Rationale

##### 1. Description of the model

When a variable y(t) exhibits a periodic pattern (e.g. circadian rhythms), its temporal evolution can be modeled as a sinusoidal oscillation around a mean level. The cosinor method estimates three physiologically interpretable parameters:

- **MESOR (M):** the mean level of the data (i.e. central value around which the rhythm oscillates).
- **Amplitude (A):** the magnitude of the oscillation around the mean.
- **Acrophase (ϕ):** the temporal position of the maximum within the period (i.e. time at which the process reaches its maximum)

##### 2. Fundamental sinusoidal model

The theoretical model assumed by the cosinor method is:

$$y(t) = M + A\cos(\omega t - \phi) + \varepsilon(t)$$

Where:

- M is the MESOR
- $A > 0$ is the amplitude
- $\phi$ is the acrophase (phase of the maximum or peak)
- $\omega = 2\pi P$ is the angular frequency
- P is the known period, which for circadian studies is fixed to 24 hours)
- $\varepsilon(t)$ is an error term

The maximum occurs when the argument of the cosine equals zero:

$\omega t - \phi = 0 \Rightarrow t = \phi / \omega$

Thus, $\phi / \omega$ corresponds to the time of the peak.

##### 3. Linearization of the model

The theoretical model assumed by the cosinor method is:

Direct estimation of A and $\phi$ is nonlinear. To enable estimation via linear regression, we exploit the trigonometric identity:

$$\cos(\omega t - \phi) = \cos(\omega t)\cos\phi + \sin(\omega t)\sin\phi$$

Substituting into the previous model yields:

$$y(t) = M + A\cos\phi \cdot \cos(\omega t) + A\sin\phi \cdot \sin(\omega t) + \varepsilon(t)$$

We then define the linear parameters:

$$\beta_0 = M, \qquad \beta_c = A\cos\phi, \qquad \beta_s = -A\sin\phi$$

With this convention, the model becomes:

$$y(t) = \beta_0 + \beta_c\cos(\omega t) - \beta_s\sin(\omega t) + \varepsilon(t)$$

This representation is linear in the parameters $\beta_0$, $\beta_c$, $\beta_s$, which allows estimation via ordinary least square (OLS).

#### 4. Recovery of physiological parameters

Once $\beta_c$ and $\beta_s$ are estimated, the original rhythm parameters are recovered as follows:

##### a. Amplitude

$$A = \sqrt{\beta c2 + \beta s2}$$

This corresponds to the Euclidean magnitude of the cosine-sine components

##### b. Phase of the rhythm

From the linear definitions:

$$\beta_c = A \cos \phi, \qquad \beta_s = -A \sin \phi$$

the acrophase is obtained by:

$$\phi = \arctan 2(-\beta_s, \beta_c)$$

The function arctan 2 preserves the correct quadrant and ensures a stable estimation.

##### c. Acrophase in time units

If the period is P (24 hours), the temporal acrophase is:

$$acrophase = \frac{P \cdot \phi}{2\pi} P$$

This directly corresponds to the time of the maximum within the cycle.

#### 5. Final equivalent model

After recovering M, A, $\phi$, the rhythm can be expressed as:

$$y(t) = M + A\cos(\omega t - \phi),$$

or equivalently:

$$y(t) = M + A\cos(\omega(t - \text{acrophase}))$$

where the acrophase indicates the peak time within the period.

#### 6. Test of rhythmicity

To assess whether a true periodic rhythm exists, the following null hypothesis is tested:

$$H_0 : \beta_c = 0 \quad \text{and} \quad \beta_s = 0$$

Under $H_0$, the model reduces to:

$$y(t) = \beta_0 + \varepsilon(t)$$

i.e. a constant mean with no oscillatory component.

A joint F-test with two linear restrictions compares: the reduced model (no rhythmic components) against the full model (with cosine and sine terms). If $H_0$ is rejected, the rhythm is considered statistically significant.

#### 7. Justification of the method

The cosinor analysis was selected as it is a robust method when the period is fixed (24h), it provides formal statistical inference, along with physiologically interpretable parameters, and it's widely established in chronobiology and circadian research.

#### Supplementary Figures

**Figure S1.** Diurnal rhythmicity of *c-Fos, Arc, Nr4a1, Dusp1, Dusp5, Dusp6*, in the male (A) and female (B) mouse hippocampus.

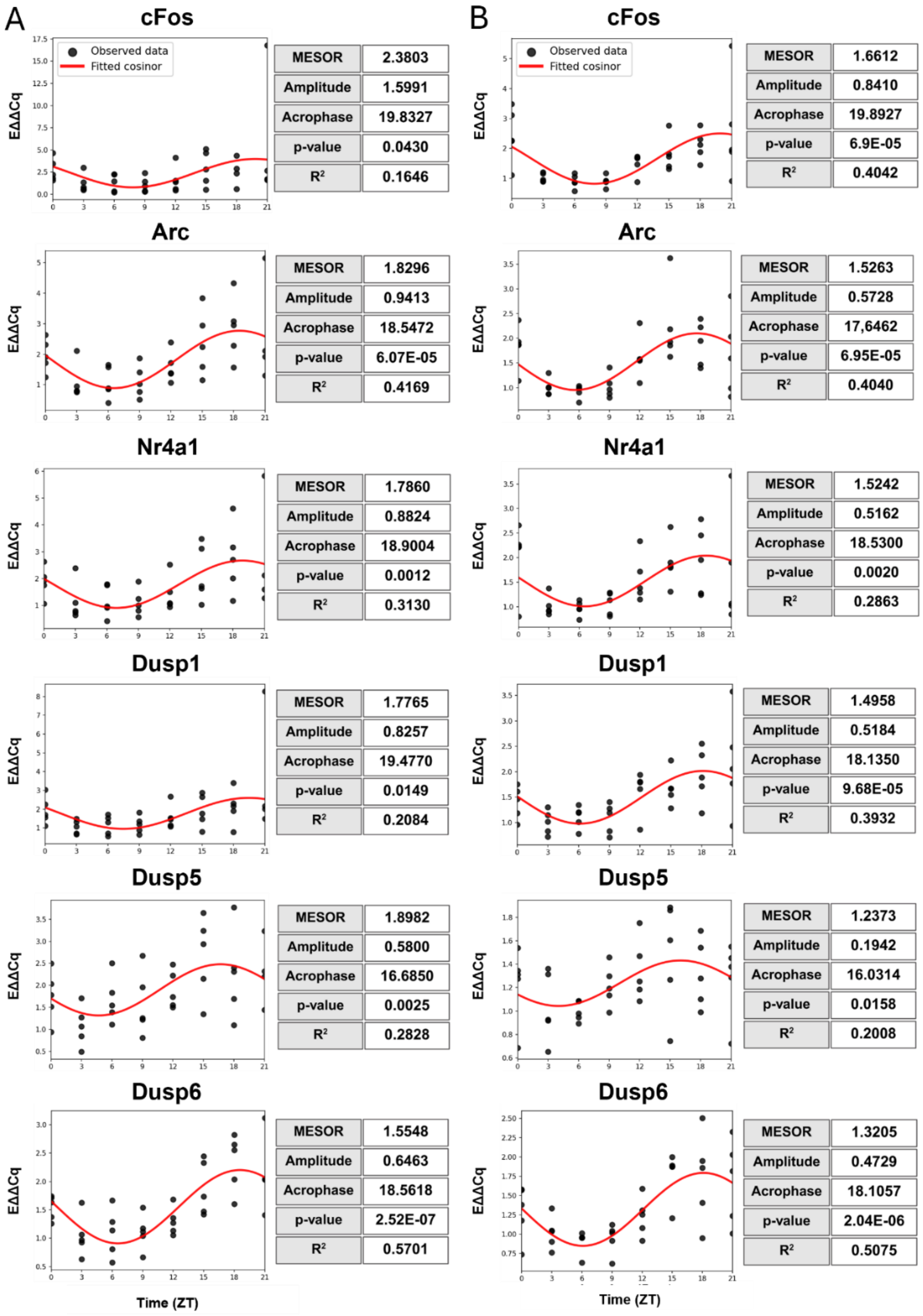


**Fig S1.** Graphs depict relative gene expression levels and fitted cosinor curve across an entire 24-hour cycle in 3-hours intervals, alongside cosinor analysis main output parameters. Dots correspond to observed individual data points and red line represents the fitted curve. Gene expression was quantified by RT-qPCR. Y-axis represents relative expression levels (EAACq) normalized to the geometric mean of *Gapdh* and *Bactin*. X-axis indicates circadian time-points (ZT) of sample collection. p-values ≤ 0.05 are considered statistically significant. Full cosinor output parameters are provided in **Supplementary Information File 4.**

**Figure S2.** Diurnal rhythmicity of *c-Fos, Arc, Nr4a1, Dusp1, Dusp5, Dusp6*, in the male (A) and female (B) mouse prefrontal cortex.

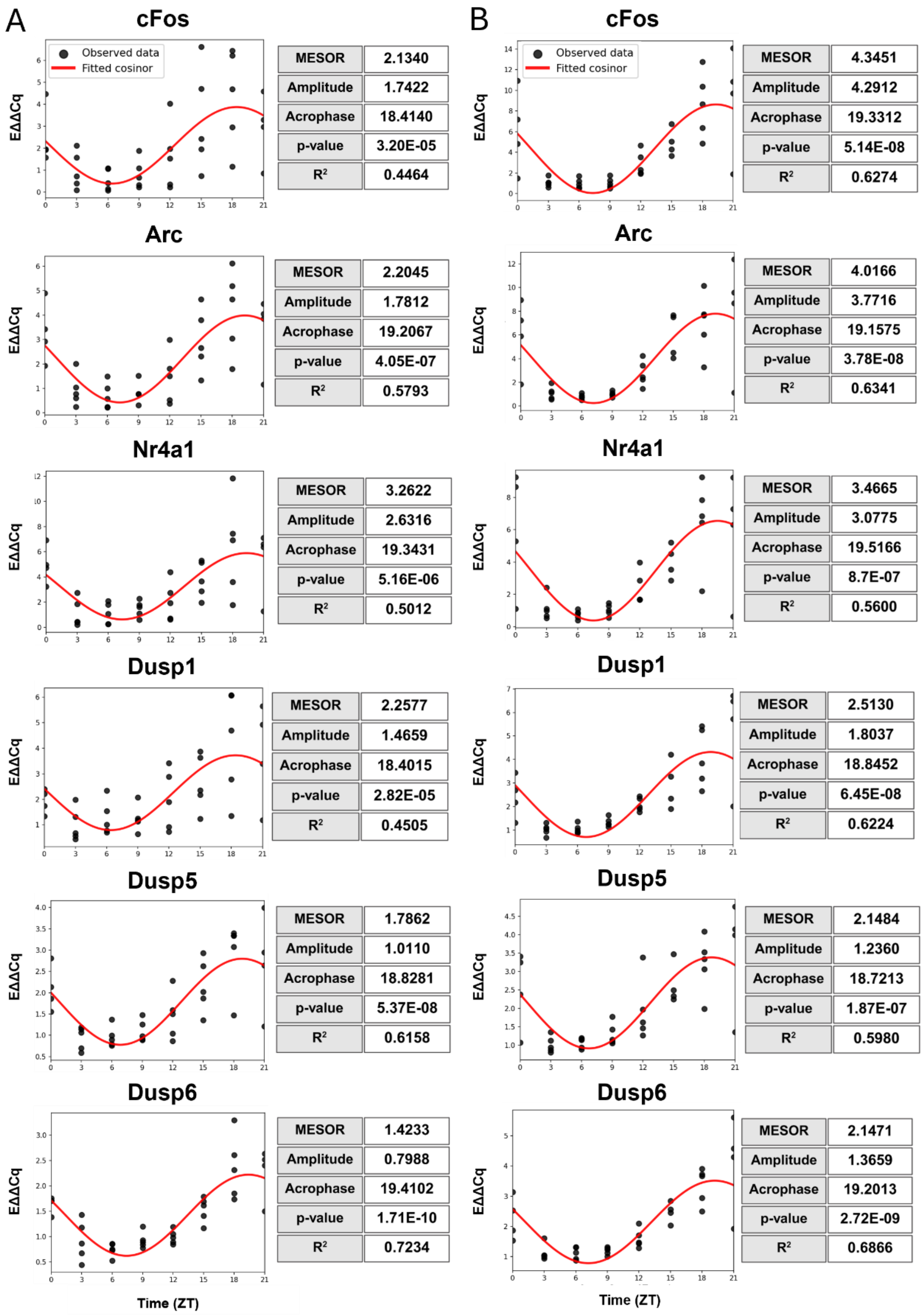


**Fig S2.** Graphs depict relative gene expression levels and fitted cosinor curve across an entire 24-hour cycle in 3-hours intervals, alongside cosinor analysis main output parameters. Dots correspond to observed individual data points and red line represents the fitted curve. Gene expression was quantified by RT-qPCR. Y-axis represents relative expression levels (EAACq) normalized to *Rplp0*. X-axis indicates circadian time-points (ZT) of sample collection. p-values ≤ 0.05 are considered statistically significant. Full cosinor output parameters are provided in **Supplementary Information File 4.**

**Figure S3.** TrkB, GSK3β and ERK2 phosphorylation patterns throughout the 24-hour cycle in the hippocampus of male (A, C, E) and female mice (B, D, F).

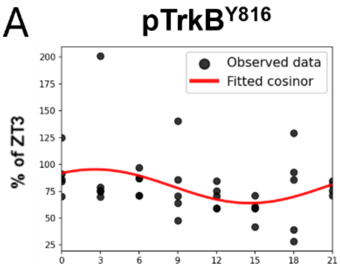


| | |
|---|---|
| MESOR | 79.6464 |
| Amplitude | 15.7050 |
| Acrophase | 2.5770 |
| p-value | 0.0576 |
| $R^2$ | 0.1467 |

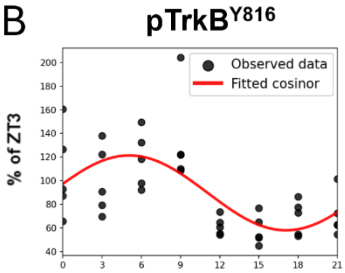


| | |
|---|---|
| MESOR | 89.6765 |
| Amplitude | 31.6731 |
| Acrophase | 5.0778 |
| p-value | 6.04E-05 |
| $R^2$ | 0.4085 |

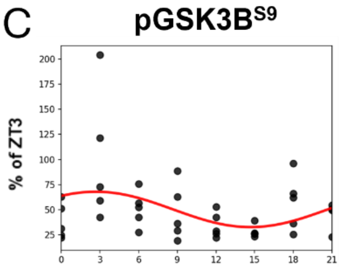


| | |
|---|---|
| MESOR | 50.3813 |
| Amplitude | 17.5998 |
| Acrophase | 2.6729 |
| p-value | 0.0667 |
| $R^2$ | 0.1396 |

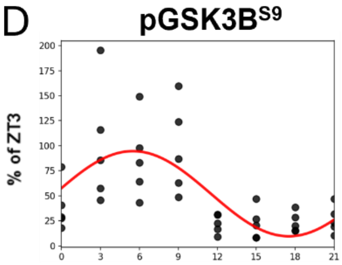


| | |
|---|---|
| MESOR | 52.0809 |
| Amplitude | 42.5380 |
| Acrophase | 5.4999 |
| p-value | 1.14E-05 |
| $R^2$ | 0.4594 |

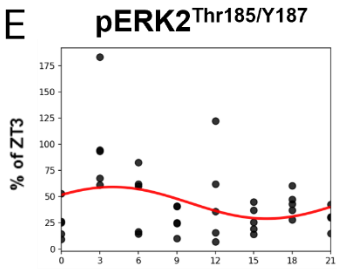


| | |
|---|---|
| MESOR | 44.0651 |
| Amplitude | 15.1185 |
| Acrophase | 3.9899 |
| p-value | 0.1535 |
| $R^2$ | 0.0989 |

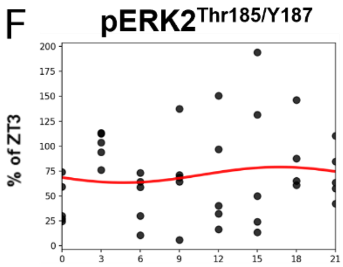


| | |
|---|---|
| MESOR | 71.1647 |
| Amplitude | 7.8421 |
| Acrophase | 16.7029 |
| p-value | 0.7420 |
| $R^2$ | 0.0164 |

**Fig S3.** Left column represents males and right column females. Graphs show the immunoblotting representation of **(A-B)** pTrkB, **(C-D)** pGSK3β, **(E-F)** pERK2, and fitted cosinor curve across an entire 24-hour cycle in 3-hours intervals, alongside cosinor analysis main output parameters. Dots correspond to observed individual data points and red line represents the fitted curve. Phosphorylation levels were quantified by western blotting. Y-axis represents phosphoproteins levels normalized against total Vinculin and compared to the average signal of ZT3, set to 100% (expressed as % of ZT3). X-axis indicates circadian time-points (ZT) of sample collection. p-values ≤ 0.05 are considered statistically significant. Full cosinor output parameters are provided in **Supplementary Information File 4.**

**Figure S4.** TrkB, GSK3β and ERK1/2 phosphorylation patterns throughout the 24-hour cycle in the prefrontal cortex of male (A, C, E, G) and female mice (B, D, F, H).

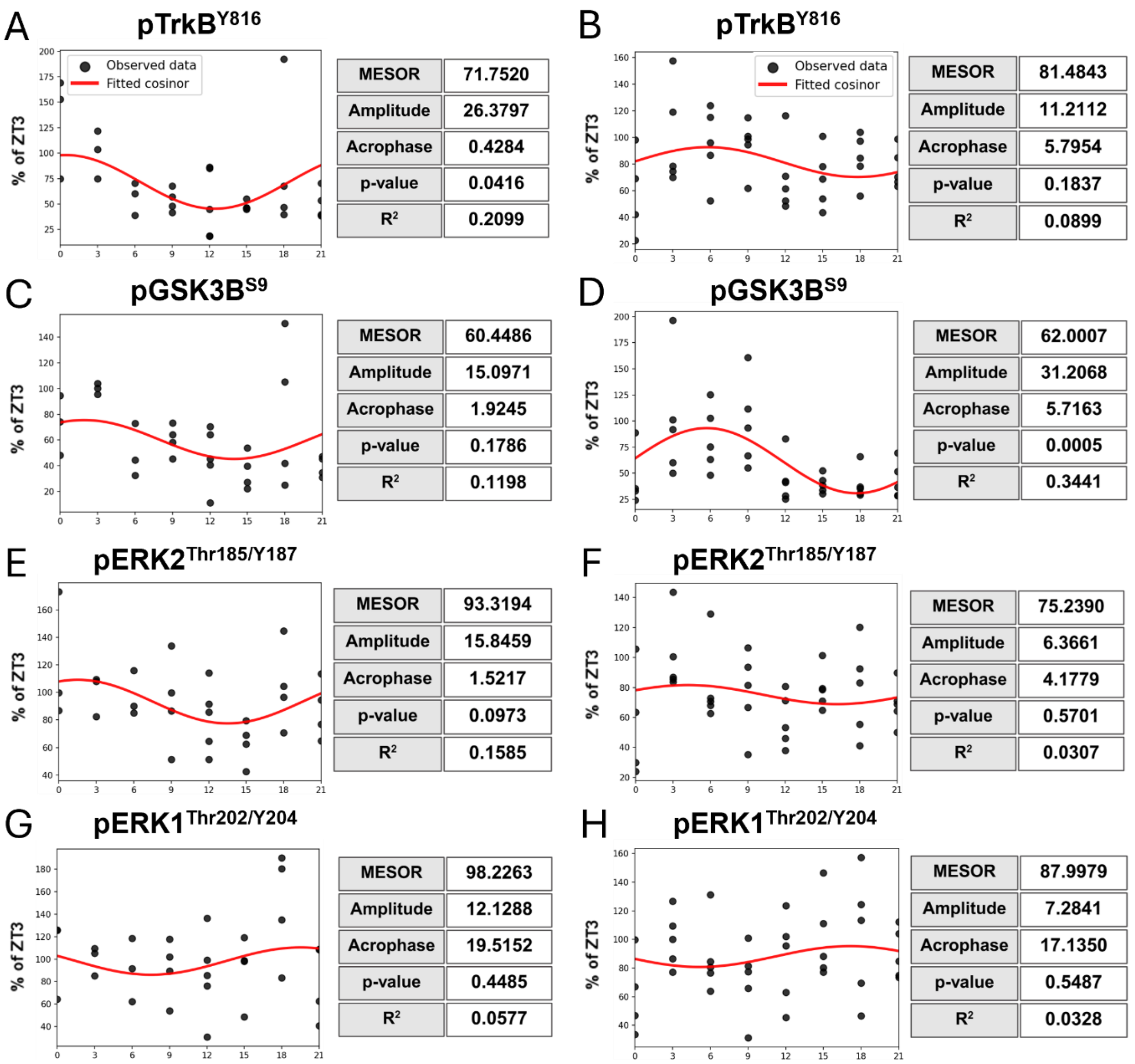


**Fig S4.** Left column represents males and right column females. Graphs show the immunoblotting representation of **(A-B)** pTrkB, **(C-D)** pGSK3β, **(E-F)** pERK2, **(G-H)** pERK1, and fitted cosinor curve across an entire 24-hour cycle in 3-hours intervals, alongside cosinor analysis main output parameters. Dots correspond to observed individual data points and red line represents the fitted curve. Phosphorylation levels were quantified by western blotting. Y-axis represents phosphoproteins levels normalized against total Vinculin and compared to the average signal of ZT3, set to 100% (expressed as % of ZT3). X-axis indicates circadian time-points (ZT) of sample collection. p-values ≤ 0.05 are considered statistically significant. Full cosinor output parameters are provided in **Supplementary Information File 4.**

**Figure S5.** Diurnal regulation of *c-Fos, Arc, Nr4a1, Dusp1, Dusp5, Dusp6*, in the male and female mouse prefrontal cortex – Additional analysis.

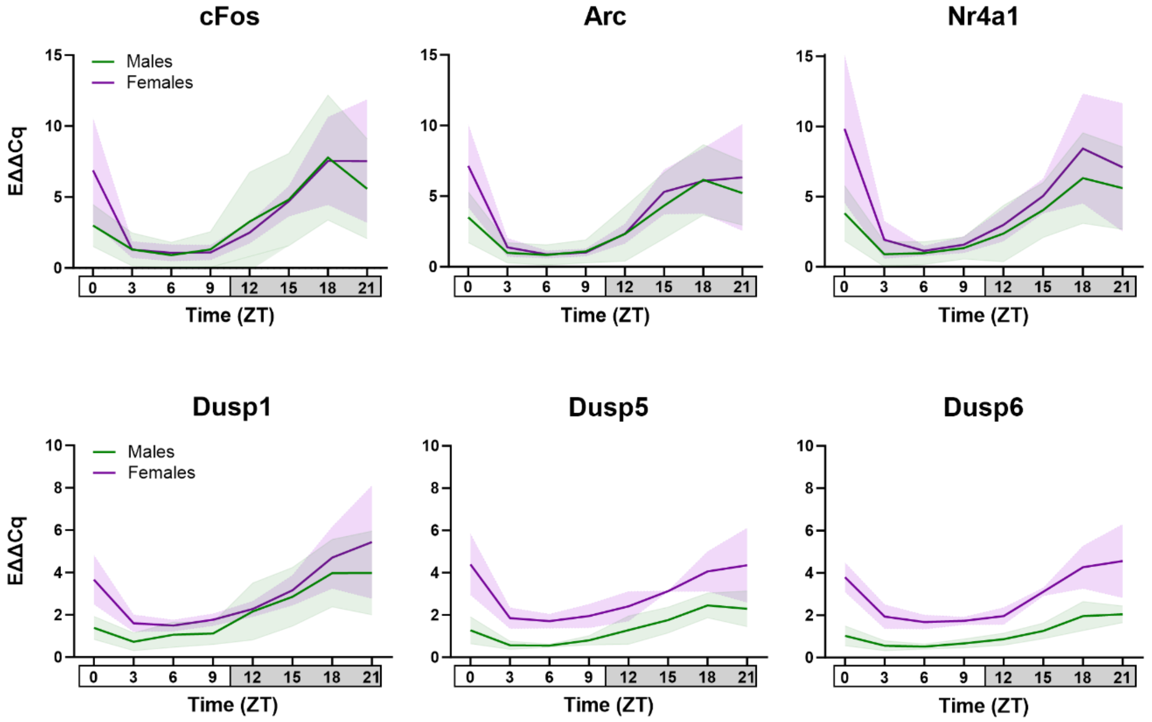


**Fig S5: Diurnal regulation of *c-Fos, Arc, Nr4a1, Dusp1, Dusp5*, *Dusp6* genes in the male (green) and female (purple) mouse prefrontal cortex.** Males are represented in green and females in purple. The graphs depict relative gene expression levels across an entire 24-hour cycle in 3-hours intervals. Solid lines represent mean values (relative to male and female ZT3 average = 1), and shaded areas indicate standard deviation (SD). Gene expression was quantified by qRT-PCR. Y-axis indicates relative gene expression (EΔΔCq) normalized to the geometric mean of Rplp0 expression from male and female samples at each circadian time-point. X-axis indicate time-points (ZT) of sample collection. n = 4-5 independent animals at each time-point. Data are presented as mean ± SD. Rhythmicity was assessed using cosinor analysis and showed statistical significance for all markers in both males and females (****$p \leq 0.0001$). Full cosinor output parameters are provided in **Supplementary Information File 4.**

#### Supplementary Tables

**Table S1.** Primers used for RT-qPCR

| Gene | Forward primer (5'→3') | Reverse primer (5'→3') |
|---|---|---|
| *Gapdh*[1] | GGTGAAGGTCGGTGTGAACGG | CATGTAGTTGAGGTCAATGAAGGG |
| *Actb*[2] *(B-actin)* | GGCTGTATTCCCCTCCATCG | CCAGTTGGTAACAATGCCATGT |
| *Rplp0*[3] | TGAGATTCGGGATATGCTGTTGG | *CGGGTCCTAGACCAGTGTTCT* |
| *Fos*[4] | CGGGTTTCAACGCCGACTA | TTGGCACTAGAGACGGACAGA |
| *Arc*[5] | AAGTGCCGAGCTGAGATGC | CGACCTGTGCAACCCTTTC |
| *Nr4a1*[6] | GGGAGTGTGCTAGAAGGACTG | AGCTTGAATACAGGGCATCTCC |
| *Dusp1*[7] | GGATATGAAGCGTTTTCGGCT | GGATTCTGCACTGTCAGGCA |
| *Dusp5*[8] | GGGGTATGAGACCTTCTACTCAC | GCGTGGTAGGCACTTCCAA |
| *Dusp6*[9] | ACAAAACTGGGCACCTTCATTC | TAGGGAAAGCGACACAGAAGTC |

[1]Karpova et al., 2010 [2]Primer bank ID: 6671509a1; [3]Primer bank ID: 254939638c1; [4]Primer bank ID: 6753894a1; [5]Primer bank ID: 9055166a1; [6]Custom made; reference sequence ID: NM_010444.2; [7]Custom-designed (Oligo 6; collaborators at the University of Strasbourg, Institute of Cellular and Integrative Neuroscience [INCI], UPR 3212, Strasbourg, France) [8]Primer bank ID: 145966802c1, [9]Custom made; reference sequence ID: NM_026268.3.

**Table S2.** Description of sample exclusions and missing values by figure and sex

**Hippocampal gene expression (Fig. 1)**: no samples were excluded for either sex, only one missing value occurred for male cFos at ZT18 due to a technical PCR failure (no Cq detected). Final sample size for males N = 39 (n = 4-5) and females N = 40 (n = 5).

**Prefrontal cortex gene expression (Fig. 2):** one male ZT0 sample and three female samples (ZT0, ZT15, ZT21) were excluded due to insufficient RNA concentration (<50 ng/µl), with one additional missing male value at ZT9 for Arc caused by a technical PCR failure (no Cq detected). Final sample size for males N = 38 (n = 4-5) and females N = 37 (n = 4-5).

**Hippocampal protein phosphorylation (Fig. 3):** no samples were excluded for either sex, although one female pERK value at ZT18 is missing due to an unquantifiable signal (dot-like artifact) resulting from abnormal protein denaturation and/or migration. Final sample size for males N = 39 (n = 4-5) and females N = 40 (n = 4-5).

**Prefrontal cortex protein phosphorylation (Fig. 4):** only 30 of the original 39 male samples were available, because several samples had to be used to troubleshoot RNA extraction issues, reducing WB sample availability across several time points (ZT0, ZT3, ZT6: −2 samples each; ZT9, ZT15, ZT18: −1 each). No female samples were excluded, however one ZT0 sample showed non-quantifiable dot-like signal across all markers. Final sample size for males N = 30 (n = 3-5) and females N = 39 (n = 4-5).

| Dataset / Figure | Sex | Planned N | Exclusions / Missing values | Reason | Final N |
|---|---|---|---|---|---|
| **Global (all datasets)** | Male | 40 | 1 excluded (ZT21) | Brain malformation | 39 |
| | Female | 40 | 0 | — | 40 |
| **Hippocampal gene expression (Fig. 1)** | Male | 39 | 1 missing value (cFos ZT18) | Technical PCR failure (no Cq detected) | 39 |
| | Female | 40 | 0 | — | 40 |
| **Prefrontal cortex gene expression (Fig. 2)** | Male | 39 | 1 excluded (ZT0)<br>1 missing val (Arc ZT9) | Low RNA conc (<40 ng/µl)<br>Technical PCR failure (no Cq) | 38 |
| | Female | 40 | 3 excluded (ZT0, ZT15, ZT21) | Low RNA conc (<40 ng/µl) | 37 |
| **Hippocampal prot phosphorylation (Fig. 3)** | Male | 39 | 0 | — | 39 |
| | Female | 40 | 1 missing value (pERK ZT18) | Non-quantifiable prot band (dot-like signal) | 40 |
| **PFC protein phosphorylation (Fig. 4)** | Male | 39 | 9 unavailable samples (ZT0, 3, 6, 9,15, 18) | Tissue used earlier for RNA extraction troubleshooting | 30 |
| | Female | 40 | 0 excluded<br>1 missing value (All markers ZT0) | —<br>Non-quantifiable prot band (dot-like signal) | 39 |

**Table S3.** Summary of statistical analyses applied in Figure 1

| Group | Sample size (n) | Shapiro-Wilk normality test (W, P value, Passed normality test?) | Statistical test (*t[df]* or *U*, p) |
|---|---|---|---|
| **FIGURE 1** | | | |
| **Male – cFos – ZT3** | 5 | 0.7999, 0.0809, Yes | Student's two-tailed unpaired *t* test |
| **Male – cFos – ZT15** | 5 | 0.9330, 0.6168, Yes | t (8) = 1.687, 0.1302 |
| **Male – Arc – ZT3** | 4* | 0.8338, 0.1778, Yes | Student's two-tailed unpaired *t* test |
| **Male – Arc – ZT18** | 5 | 0.9755, 0.9095, Yes | t (7) = 3.856, 0.0062** |
| **Male – Nr4a1 – ZT3** | 5 | 0.7492, 0.0293, No | Mann-Whitney test |
| **Male – Nr4a1 – ZT18** | 5 | 0.9848, 0.9584, Yes | 2, 0.0317* |
| **Male – Dusp1 – ZT3** | 5 | 0.9095, 0.4646, Yes | Student's two-tailed unpaired *t* test |
| **Male – Dusp1 – ZT18** | 5 | 0.9607, 0.8131, Yes | t (8) = 2.349, 0.0468* |
| **Male – Dusp5 – ZT3** | 5 | 0.9978, 0.9986, Yes | Student's two-tailed unpaired *t* test |
| **Male – Dusp5 – ZT15** | 5 | 0.9518, 0.7499, Yes | t (8) = 3.469, 0.0085** |
| **Male – Dusp6 – ZT3** | 5 | 0.9112, 0.4749, Yes | Student's two-tailed unpaired *t* test |
| **Male – Dusp6 – ZT18** | 5 | 0.9100, 0.4676, Yes | t (8) = 4.645, 0.0017** |
| **Female – cFos – ZT3** | 5 | 0.8313, 0.1422, Yes | Student's two-tailed unpaired *t* test |
| **Female – cFos – ZT18** | 5 | 0.9982, 0.9990, Yes | t (8) = 4.703, 0.0015** |
| **Female – Arc – ZT3** | 5 | 0.8192, 0.1152, Yes | Student's two-tailed unpaired *t* test |
| **Female – Arc – ZT15** | 5 | 0.7788, 0.0538, Yes | t (8) = 3.386, 0.0096** |
| **Female – Nr4a1 – ZT3** | 5 | 0.7970, 0.0766, Yes | Student's two-tailed unpaired *t* test |
| **Female – Nr4a1 – ZT18** | 5 | 0.9002, 0.4110, Yes | t (8) = 2.881, 0.0205* |
| **Female – Dusp1 – ZT3** | 5 | 0.9734, 0.8963, Yes | Student's two-tailed unpaired *t* test |
| **Female – Dusp1 – ZT18** | 5 | 0.9726, 0.8918, Yes | t (8) = 3.555, 0.0075* |
| **Female – Dusp5 – ZT6** | 5 | 0.8944, 0.3796, Yes | Student's two-tailed unpaired *t* test |
| **Female – Dusp5 – ZT15** | 5 | 0.9476, 0.7202, Yes | t (8) = 2.186, 0.0603 |
| **Female – Dusp6 – ZT6** | 4* | 0.6896, 0.0087, No | Mann-Whitney test |
| **Female – Dusp6 – ZT15** | 5 | 0.6978, 0.0091, No | 0, 0.0159* |

*1 value identified as an outlier according to Grubb's test (α = 0.01)

**Table S4.** Summary of statistical analyses applied in Figure 2

| Group | Sample size (n) | Shapiro-Wilk normality test (W, P value, Passed normality test?) | Statistical test (*t[df]* or *U*, p-value) |
|---|---|---|---|
| **FIGURE 2** | | | |
| Male – cFos – ZT3 | 5 | 0.9345, 0.6272, Yes | Student's two-tailed unpaired *t* test |
| Male – cFos – ZT18 | 5 | 0.9189, 0.5226, Yes | t (8) = 3.093, 0.0148* |
| Male – Arc – ZT3 | 5 | 0.9199, 0.5296, Yes | Student's two-tailed unpaired *t* test |
| Male – Arc – ZT18 | 5 | 0.9606, 0.8120, Yes | t (8) = 3.888, 0.0046** |
| Male – Nr4a1 – ZT3 | 5 | 0.8328, 0.1459, Yes | Student's two-tailed unpaired *t* test |
| Male – Nr4a1 – ZT18 | 5 | 0.9649, 0.8416, Yes | t (8) = 2.881, 0.0205* |
| Male – Dusp1 – ZT3 | 5 | 0.8718, 0.2739, Yes | Student's two-tailed unpaired *t* test |
| Male – Dusp1 – ZT18 | 5 | 0.8915, 0.3647, Yes | t (8) = 3.276, 0.0113* |
| Male – Dusp5 – ZT3 | 5 | 0.8497, 0.1937, Yes | Student's two-tailed unpaired *t* test |
| Male – Dusp5 – ZT18 | 4* | 0.7986, 0.0997, Yes | t (7) = 15.50, <0.0001**** |
| Male – Dusp6 – ZT3 | 5 | 0.9780, 0.9239, Yes | Student's two-tailed unpaired *t* test |
| Male – Dusp6 – ZT18 | 5 | 0.9360, 0.6381, Yes | t (8) = 4.350, 0.0024** |
| Female – cFos – ZT3 | 5 | 0.9064, 0.4464, Yes | Student's two-tailed unpaired *t* test |
| Female – cFos – ZT18 | 5 | 0.9807, 0.9381, Yes | t (8) = 5.307, 0.0007*** |
| Female – Arc – ZT3 | 5 | 0.9290, 0.5895, Yes | Student's two-tailed unpaired *t* test |
| Female – Arc – ZT18 | 5 | 0.9666, 0.8530, Yes | t (8) = 5.050, 0.0010*** |
| Female – Nr4a1 – ZT3 | 5 | 0.8203, 0.1174, Yes | Student's two-tailed unpaired *t* test |
| Female – Nr4a1 – ZT18 | 5 | 0.9019, 0.4204, Yes | t (8) = 4.368, 0.0024** |
| Female – Dusp1 – ZT3 | 5 | 0.9783, 0.9256, Yes | Student's two-tailed unpaired *t* test |
| Female – Dusp1 – ZT18 | 5 | 0.9018, 0.4199, Yes | t (8) = 5.448, 0.0006*** |
| Female – Dusp5 – ZT3 | 5 | 0.9185, 0.5203, Yes | Student's two-tailed unpaired *t* test |
| Female – Dusp5 – ZT18 | 5 | 0.9449, 0.7008, Yes | t (8) = 6.037, 0.0003*** |
| Female – Dusp6 – ZT3 | 4* | 0.9478, 0.7024, Yes | Student's two-tailed unpaired *t* test |
| Female – Dusp6 – ZT18 | 5 | 0.8761, 0.2921, Yes | t (7) = 7.682, 0.0001*** |

*1 value identified as an outlier according to Grubb's test (α = 0.01)

**Table S5.** Summary of statistical analyses applied in Figure 3

| Group | Sample size (n) | Shapiro-Wilk normality test (W, P value, Passed normality test?) | Statistical test (*t[df]* or *U*, p-value) |
|---|---|---|---|
| **FIGURE 3** | | | |
| Male – pTrkB – ZT3 | 4* | 0.9584, 0.7691, Yes | Student's two-tailed unpaired *t* test |
| Male – pTrkB – ZT15 | 5 | 0.8889, 0.3514, Yes | t (7) = 2.957, 0.0212* |
| Male – pGSK3B – ZT3 | 5 | 0.8820, 0.3183, Yes | Mann-Whitney test |
| Male – pGSK3B – ZT15 | 5 | 0.7490, 0.0291, No | 0, 0.0079** |
| Male – pERK2 – ZT3 | 5 | 0.8021, 0.0843, Yes | Student's two-tailed unpaired *t* test |
| Male – pERK2– ZT15 | 5 | 0.9572, 0.7885, Yes | t (8) = 3.180, 0.0130* |
| Female – pTrkB – ZT6 | 5 | 0.9504, 0.7398, Yes | Student's two-tailed unpaired *t* test |
| Female – pTrkB – ZT15 | 5 | 0.9282, 0.5840, Yes | t (8) = 4.986, 0.0011** |
| Female – pGSK3B – ZT3 | 5 | 0.9014, 0.4174, Yes | Student's two-tailed unpaired *t* test |
| Female – pGSK3B – ZT15 | 5 | 0.8922, 0.3683, Yes | t (8) = 2.808, 0.0229* |
| Female – pERK2 – ZT3 | 5 | 0.8975, 0.3964, Yes | Student's two-tailed unpaired *t* test |
| Female – pERK2– ZT15 | 5 | 0.8838, 0.3267, Yes | t (8) = 0.4896, 0.6376 |

*1 value identified as an outlier according to Grubb's test (α = 0.01)

**Table S6.** Summary of statistical analyses applied in Figure 4

| Group | Sample size (n) | Shapiro-Wilk normality test (W, P value, Passed normality test?) | Statistical test (*t[df]* or *U*, p-value) |
|---|---|---|---|
| **FIGURE 4** | | | |
| Male – pTrkB – ZT3 | 3 | 0.9839, 0.7567, Yes | Student's two-tailed unpaired *t* test |
| Male – pTrkB – ZT15 | 4 | 0.7802, 0.0711, Yes | t (5) = 4.411, 0.007** |
| Male – pGSK3B – ZT3 | 3 | 0.9957, 0.8742, Yes | Student's two-tailed unpaired *t* test |
| Male – pGSK3B – ZT15 | 4 | 0.9443, 0.6807, Yes | t (5) = 7.483, 0.0007*** |
| Male – pERK2 – ZT3 | 3 | 0.7953, 0.1033, Yes | Student's two-tailed unpaired *t* test |
| Male – pERK2– ZT15 | 4 | 0.9651, 0.8109, Yes | t (5) = 3.109, 0.0266* |
| Male – pERK1 – ZT3 | 3 | 0.8781, 0.3189, Yes | Student's two-tailed unpaired *t* test |
| Male – pERK1– ZT15 | 4 | 0.8751, 0.3181, Yes | t (5) = 0.4660, 0.6608 |
| Female – pTrkB – ZT3 | 5 | 0.8353, 0.1524, Yes | Student's two-tailed unpaired *t* test |
| Female – pTrkB – ZT15 | 5 | 0.9778, 0.9223, Yes | t (8) = 1.893, 0.0951 |
| Female – pGSK3B – ZT6 | 5 | 0.9609, 0.8141, Yes | Student's two-tailed unpaired *t* test |
| Female – pGSK3B – ZT15 | 5 | 0.9643, 0. 8377, Yes | t (8) = 3.014, 0.0167* |
| Female – pERK2 – ZT3 | 5 | 0.7408, 0.0244, No | Mann-Whitney test |
| Female – pERK2– ZT15 | 5 | 0.9030, 0.4268, Yes | 4, 0.0952 |
| Female – pERK2 – ZT3 | 5 | 0.9827, 0.9485, Yes | Student's two-tailed unpaired *t* test |
| Female – pERK2– ZT15 | 5 | 0.8582, 0.2220, Yes | t (8) = 0.0379, 0.9707 |

#### Full blots

**Figure 3A:** pTrkB Gel 1-4 (Males)

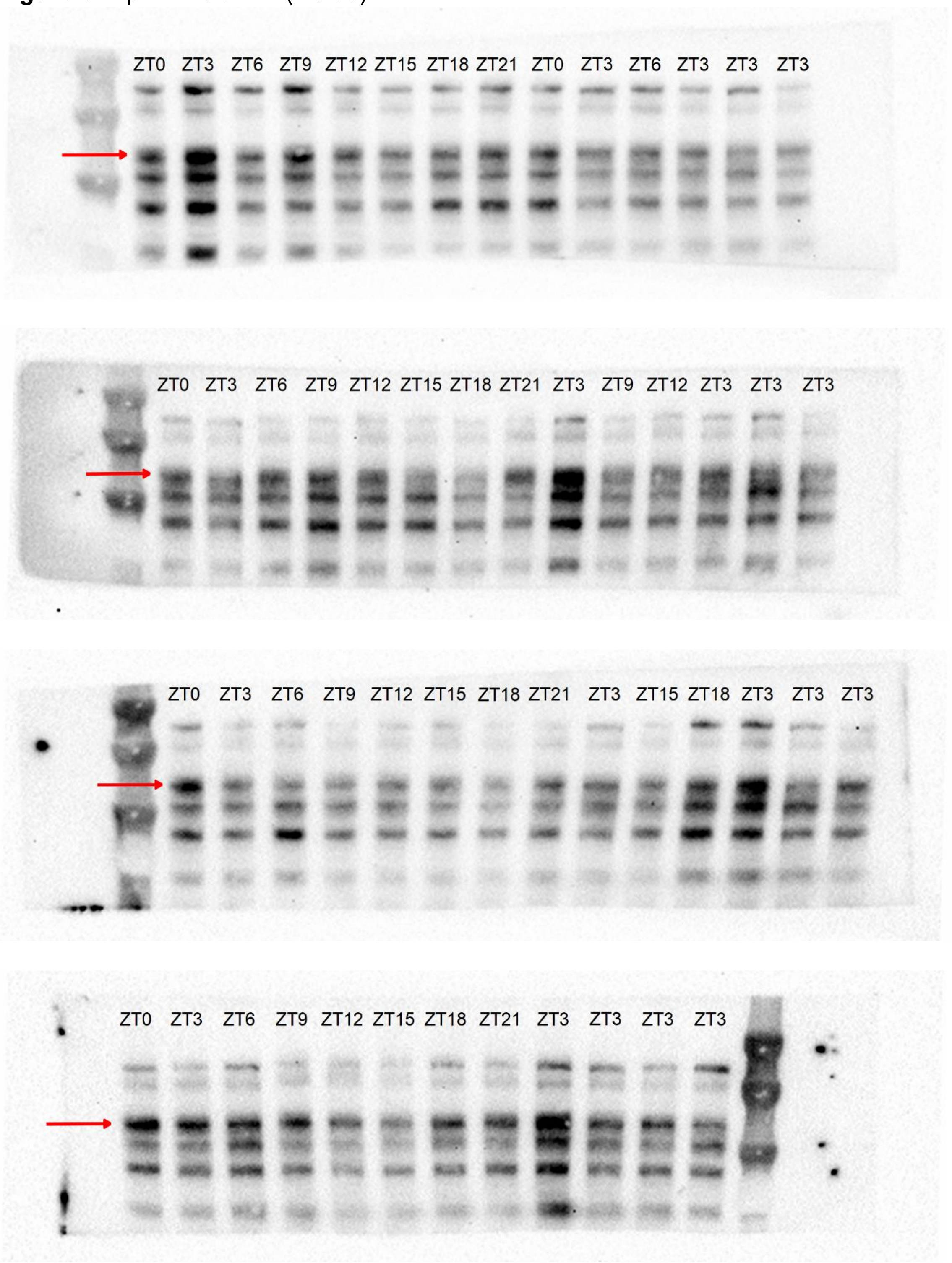

**Figure 3B:** pTrkB Gel 1-4 (Females)

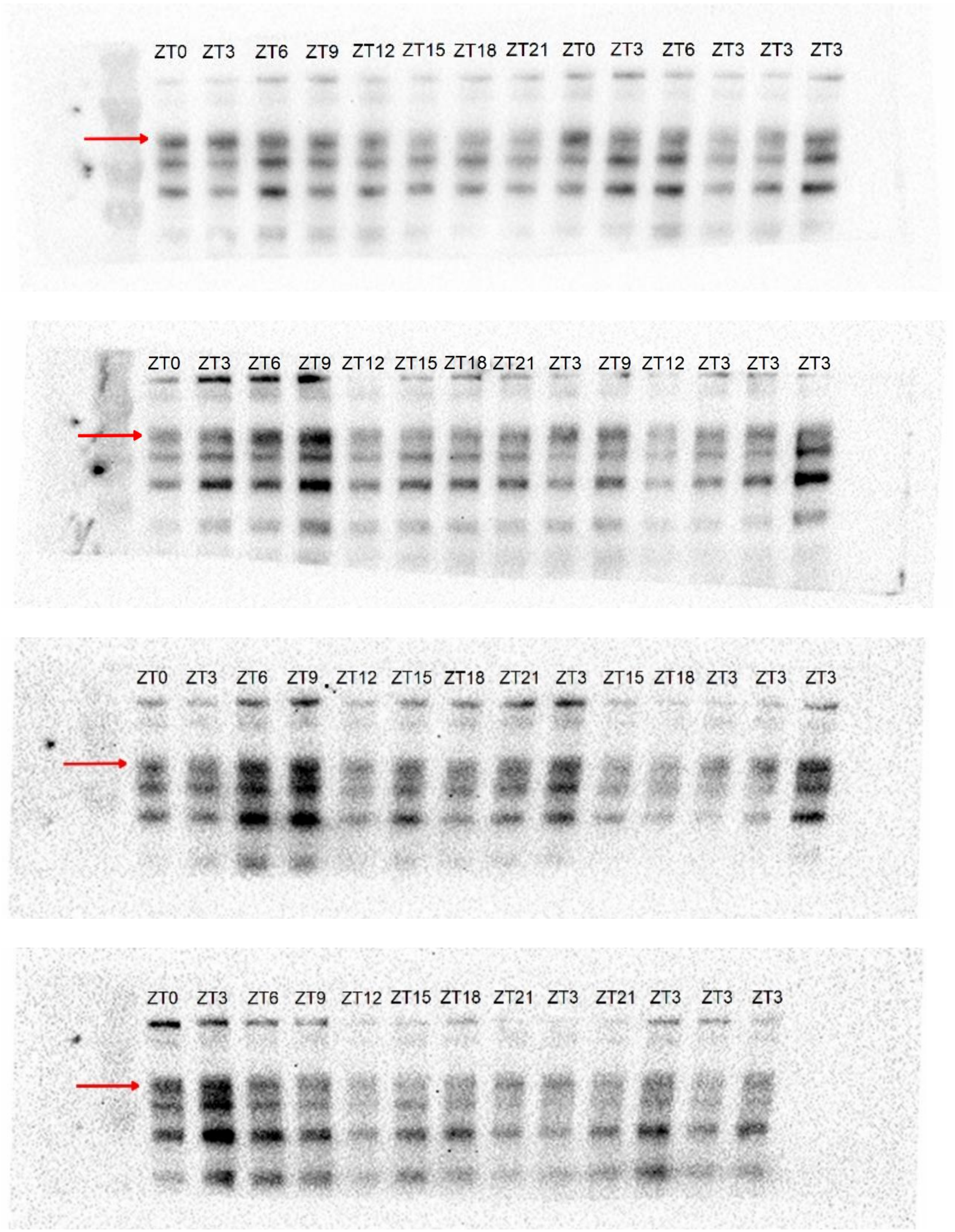

**Figure 3C:** pGSK3β Gel 1-4 (Males)

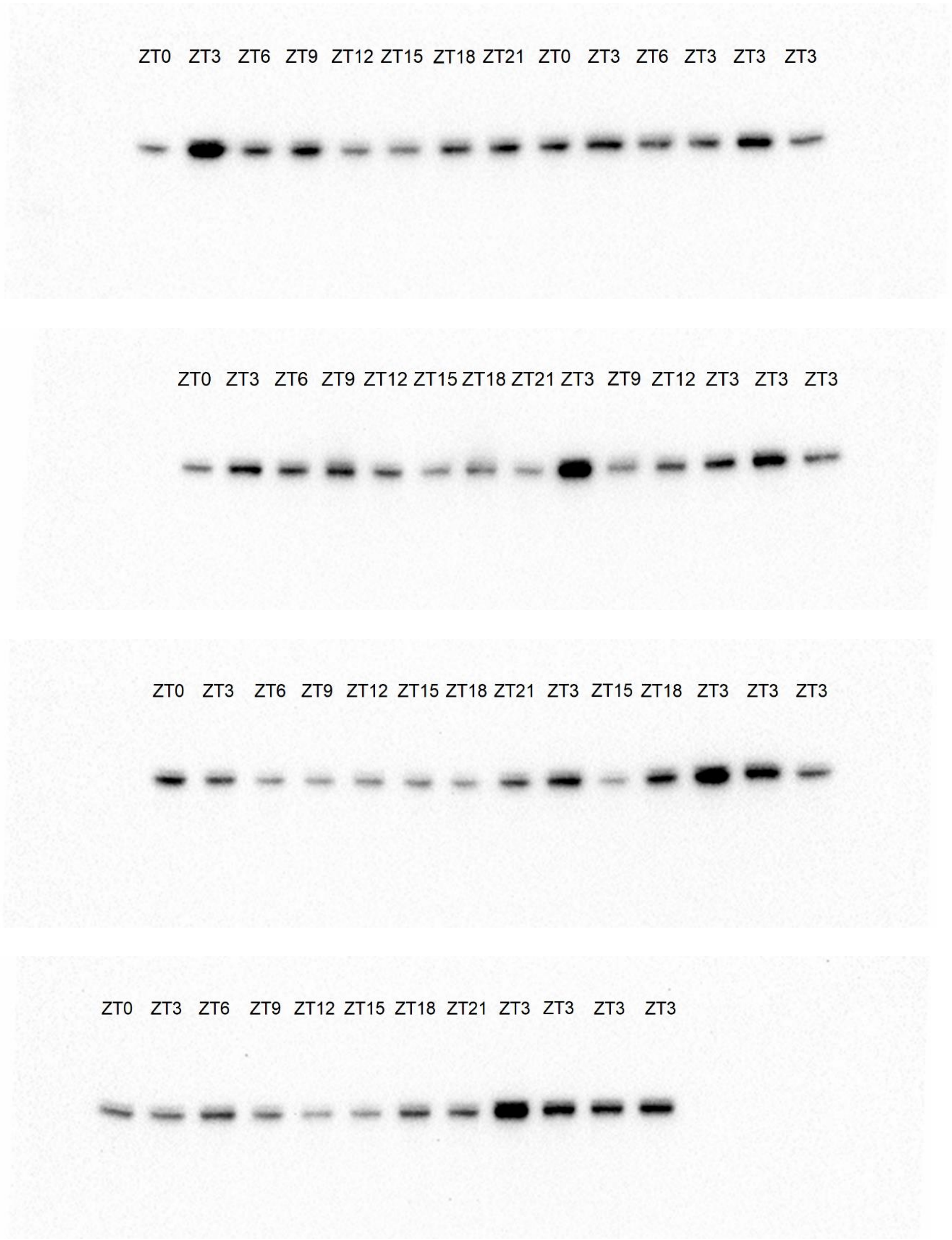

**Figure 3D:** pGSK3β Gel 1-4 (Females)

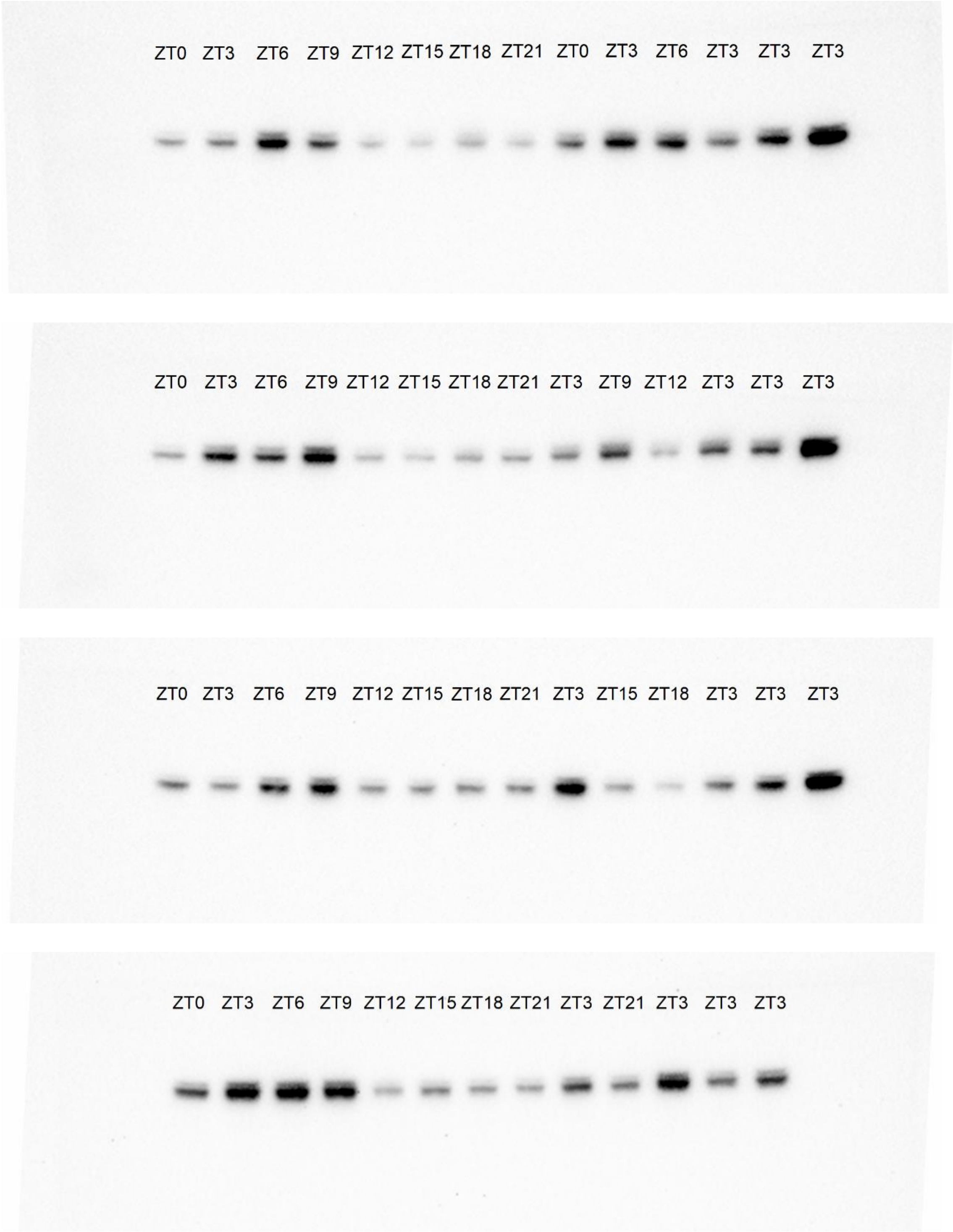

**Figure 3E:** pERK2 Gel 1-4 (Males)

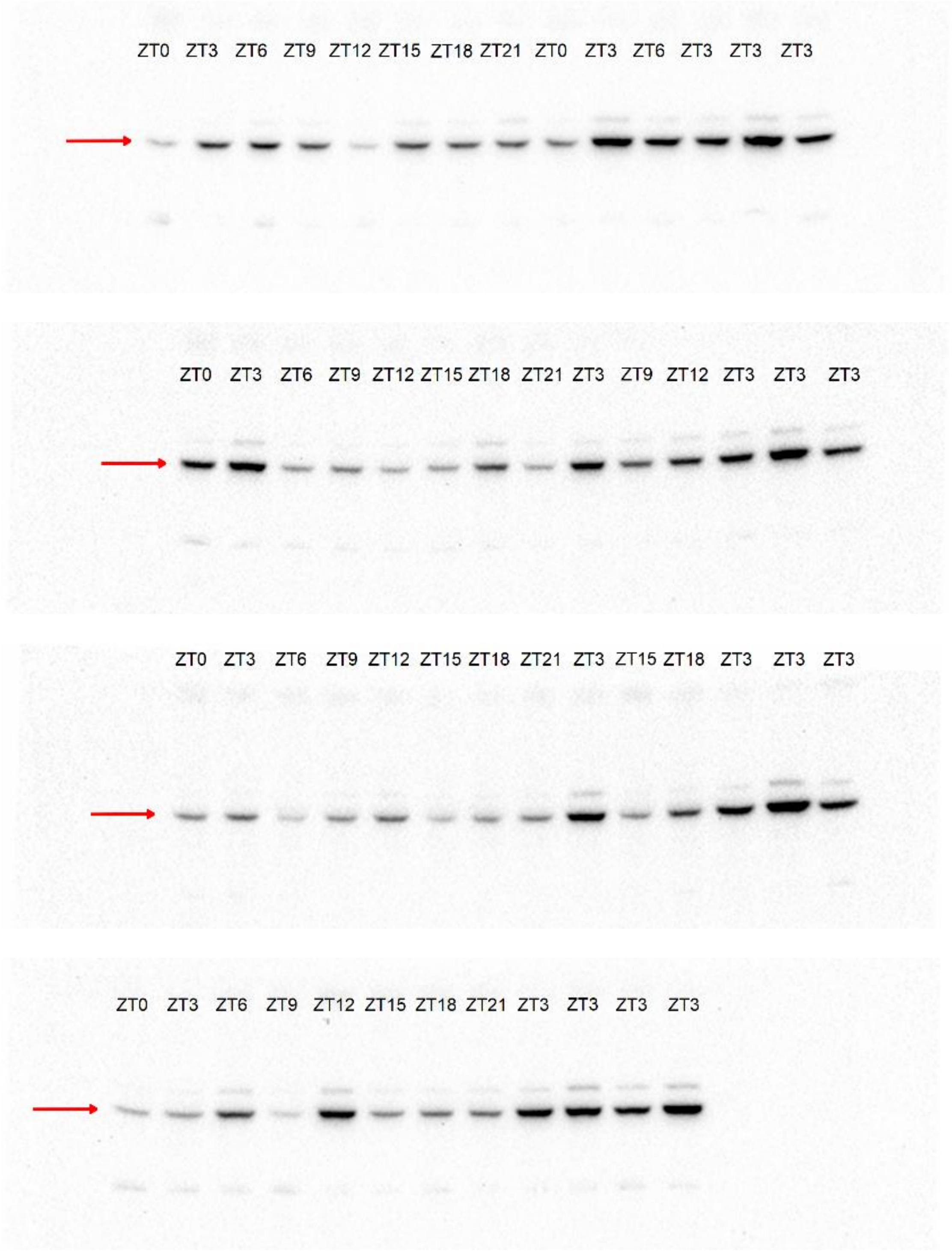

**Figure 3F:** pERK2 Gel 1-4 (Females)

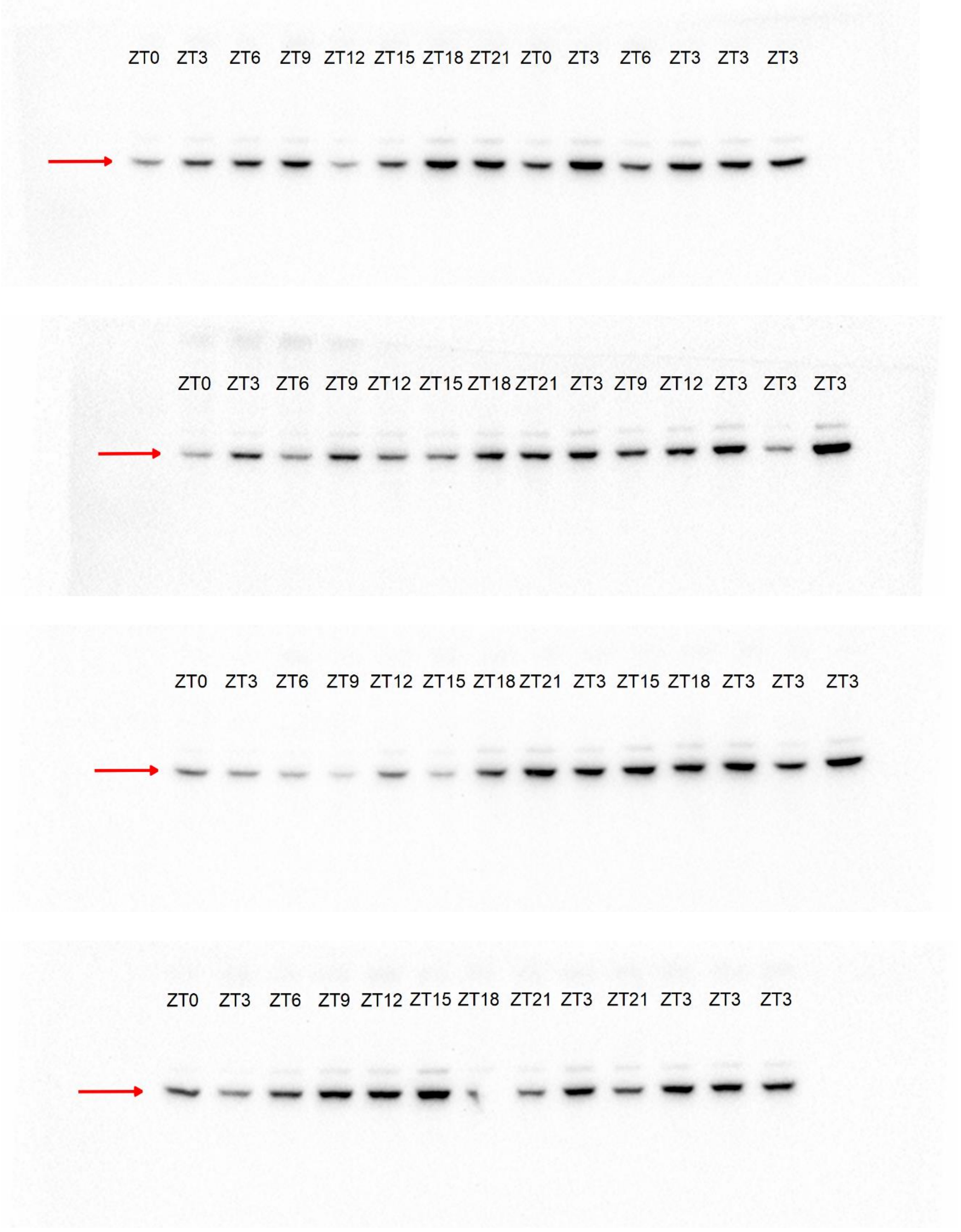

**Figure 3A, C, E:** Vinculin Gel 1-4 (Males)

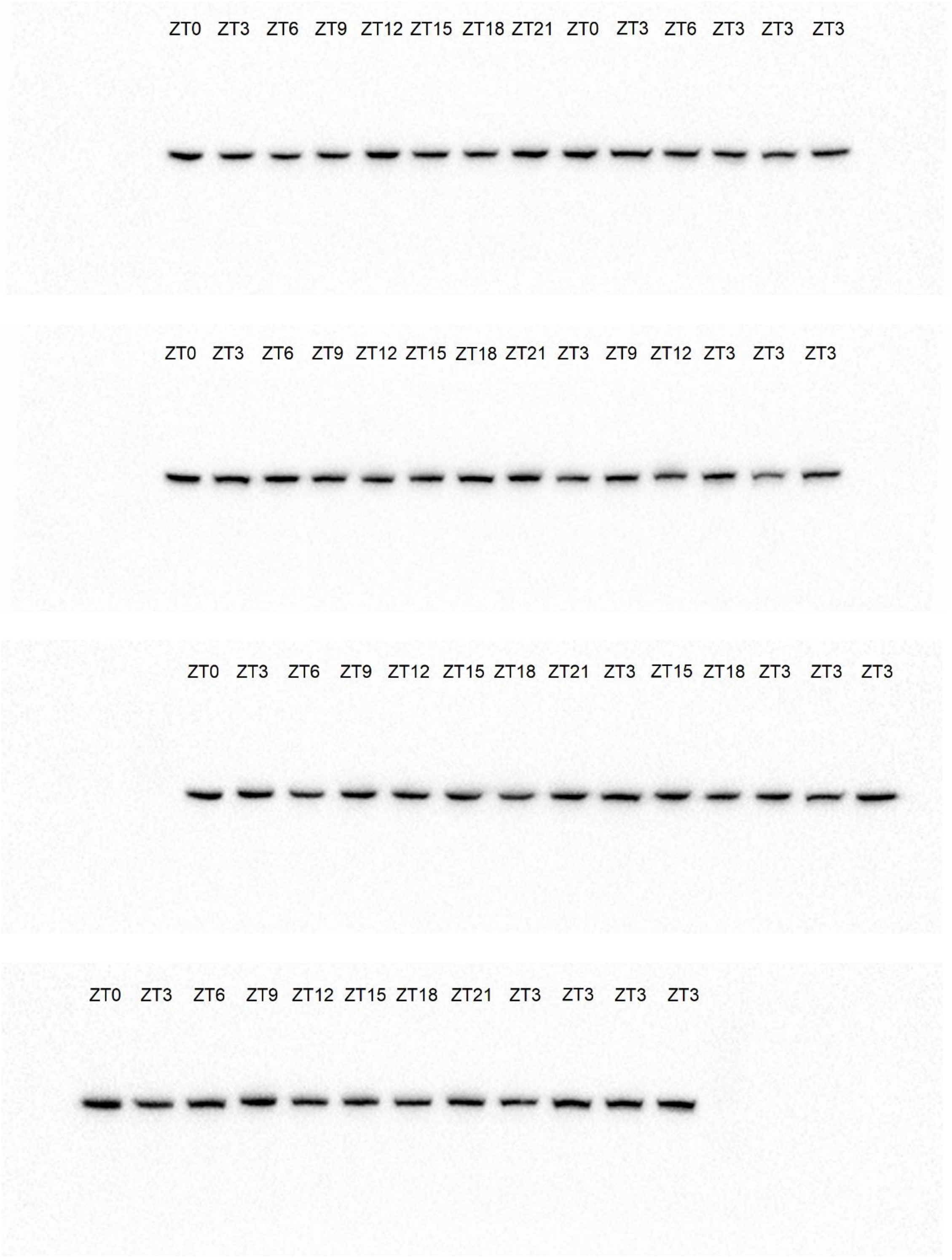

**Figure 3B, D, F:** Vinculin Gel 1-4 (Females)

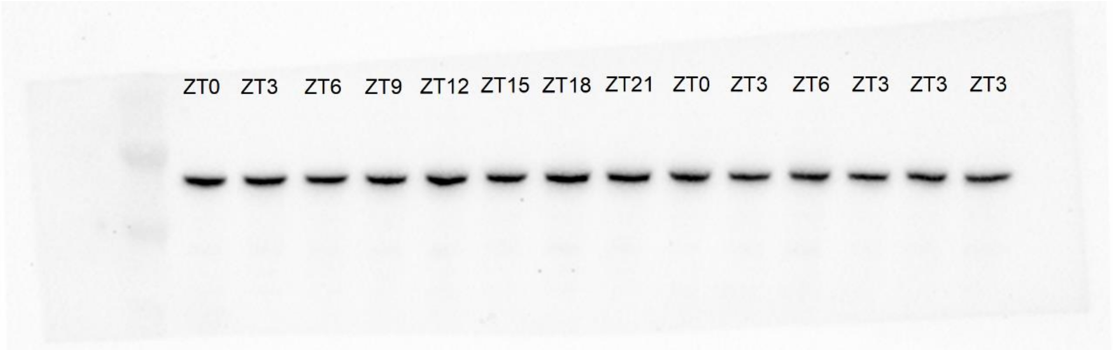


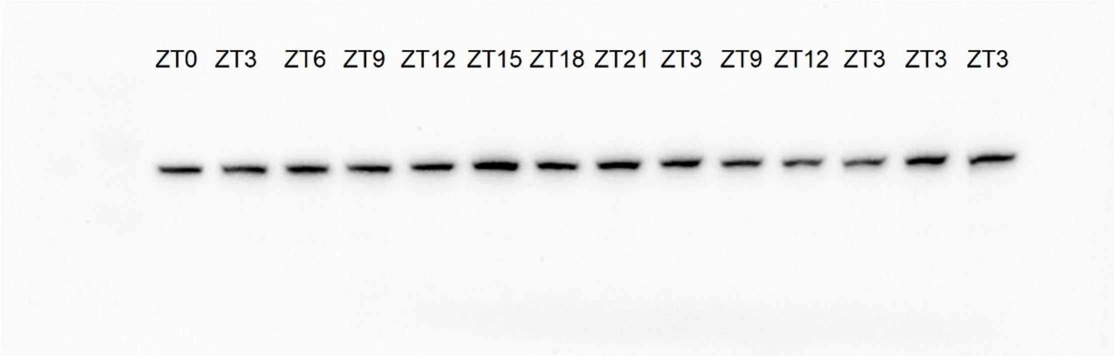


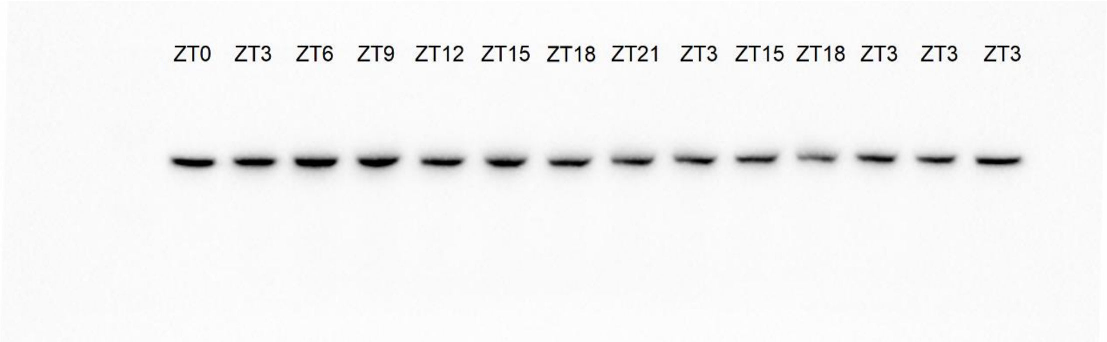


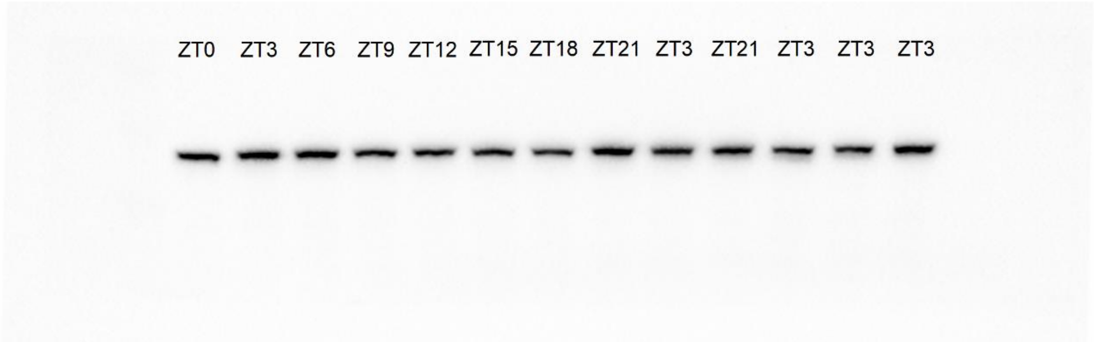

**Figure 4A:** pTrkB Gel 1-3 (Males)

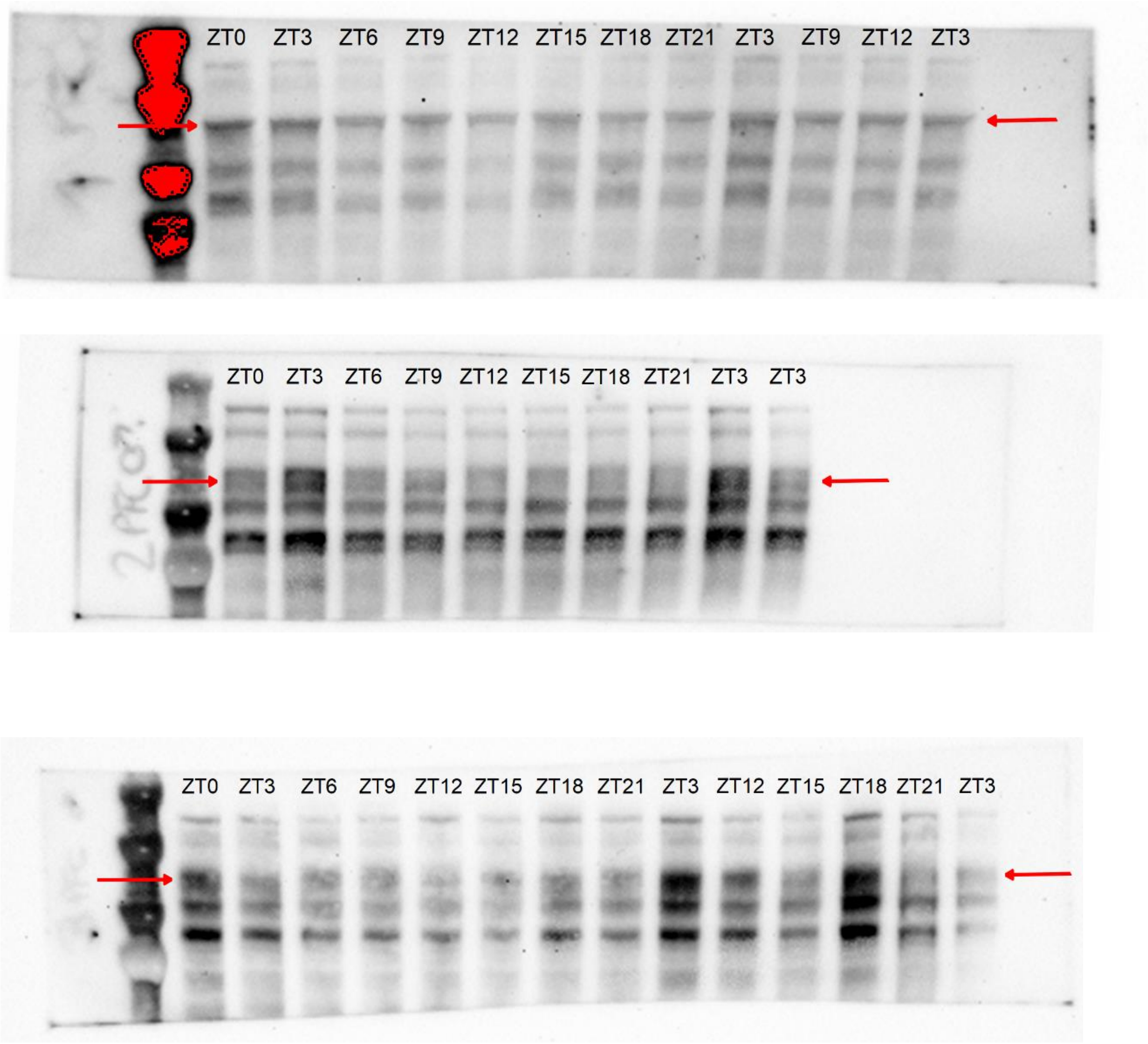

**Figure 4B:** pTrkB Gel 1-4 (Females)

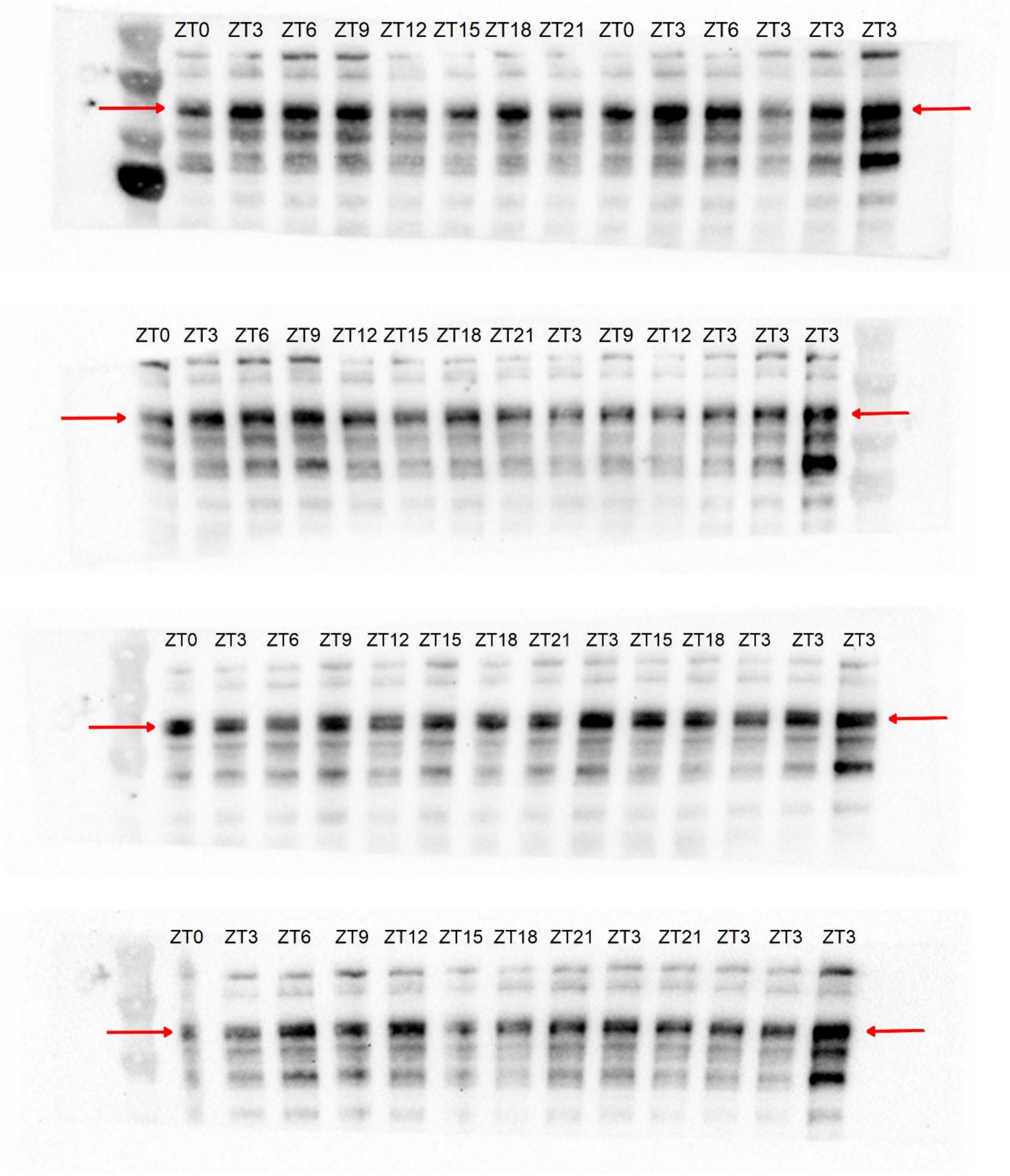

**Figure 4C:** pGSK3β Gel 1-3 (Males)

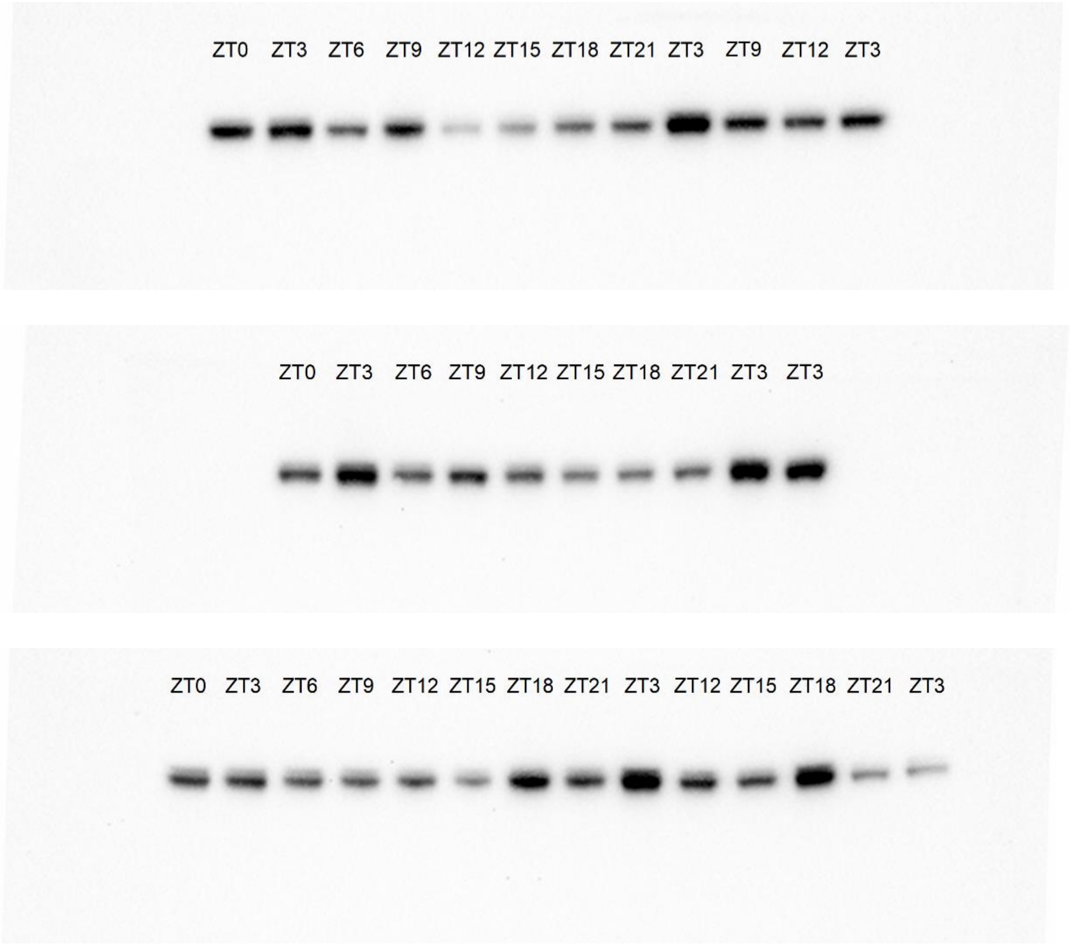

**Figure 4D:** pGSK3β Gel 1-4 (Females)

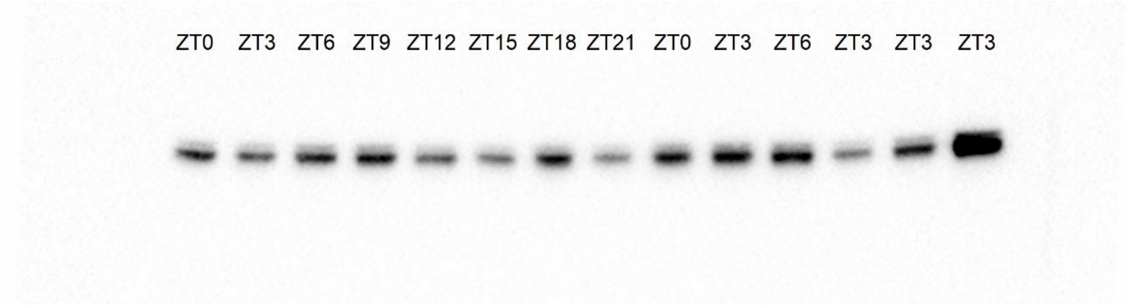


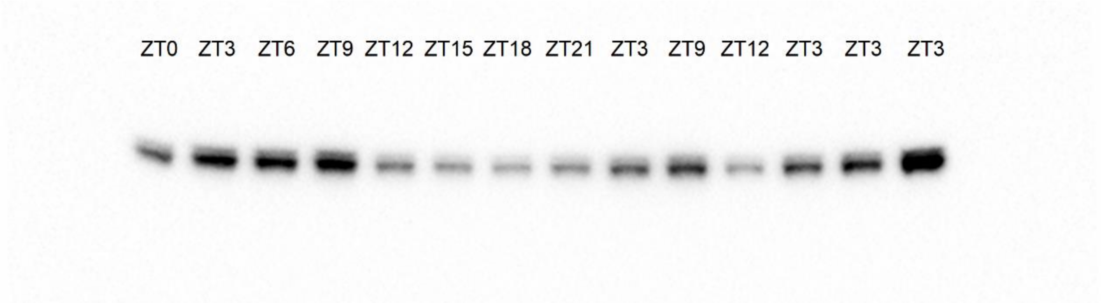


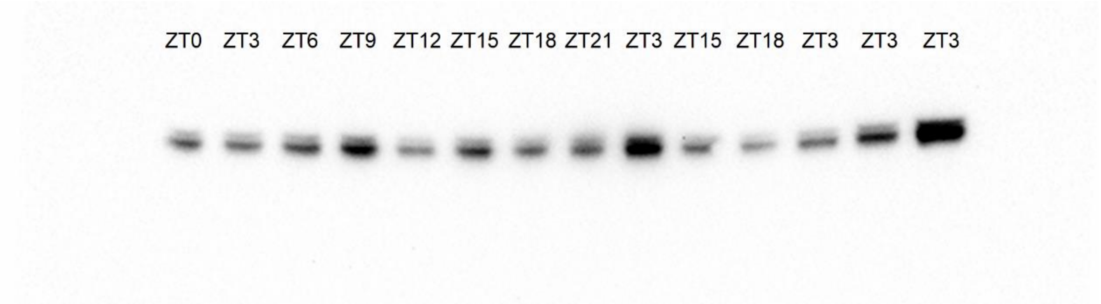


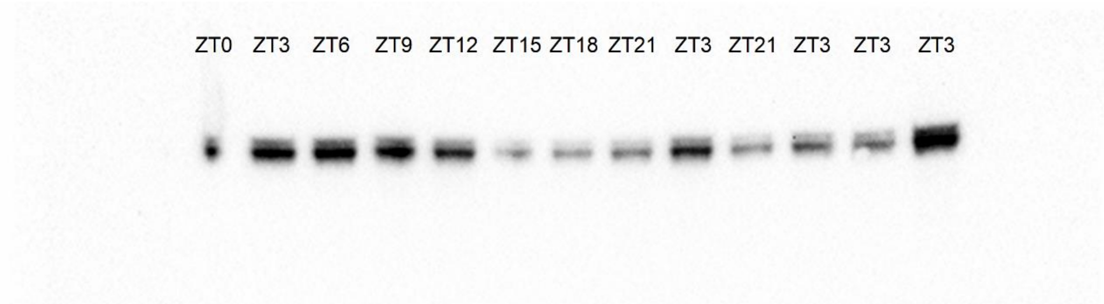

**Figure 4E, G:** pERK1/2 Gel 1-3 (Males)

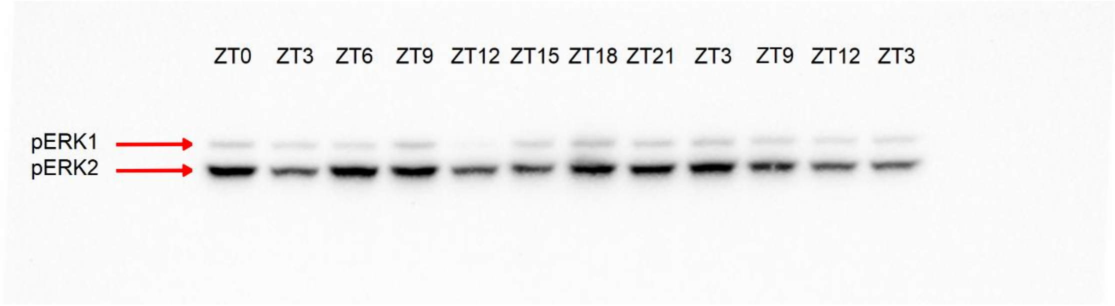


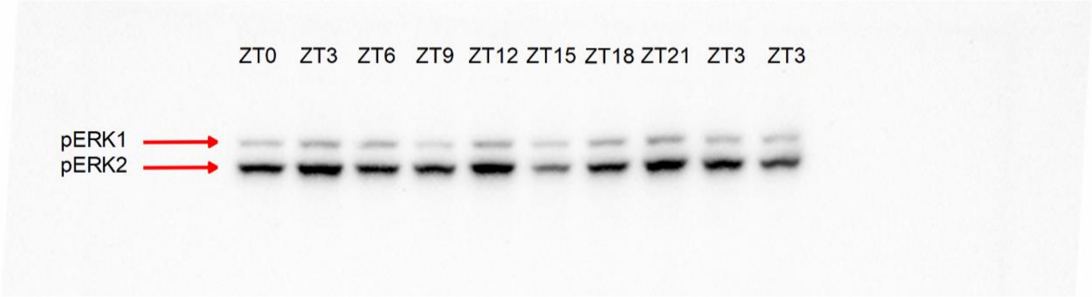


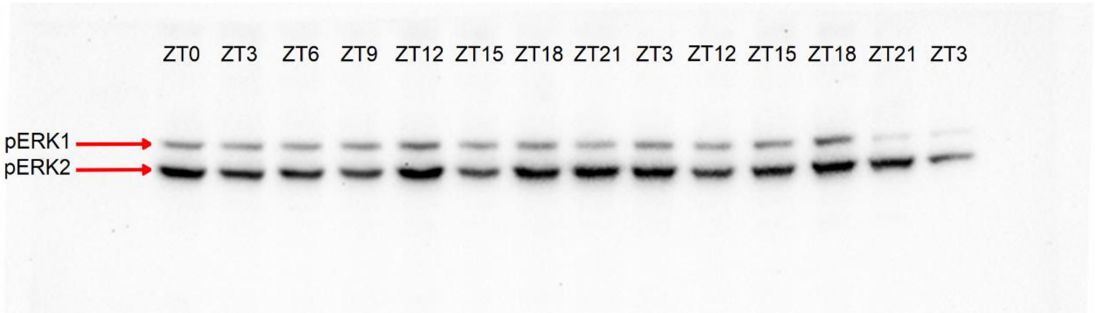

**Figure 4F, H:** pERK1/2 Gel 1-4 (Females)

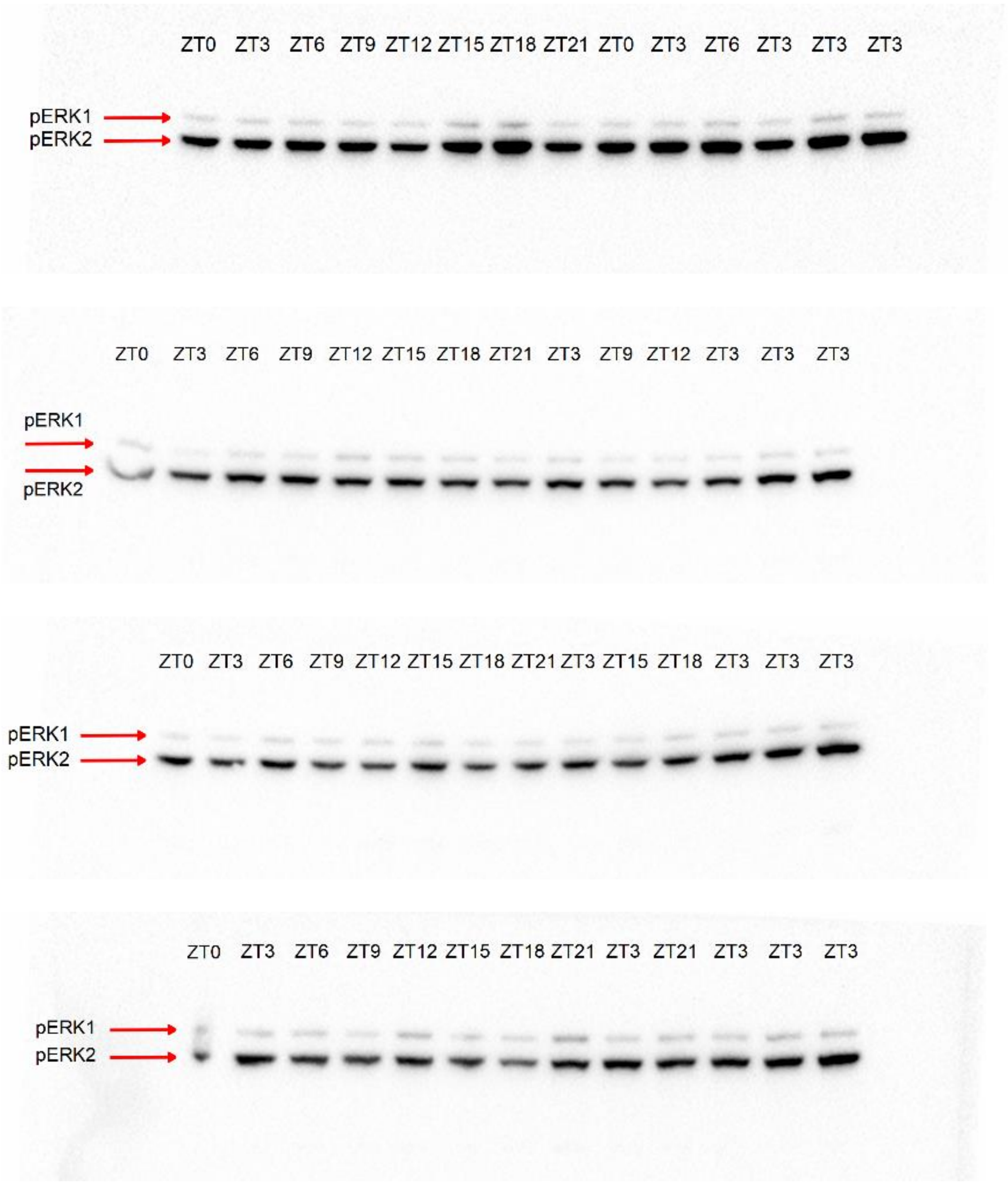

**Figure 4A, C, E, G:** Vinculin Gel 1-3 (Males)

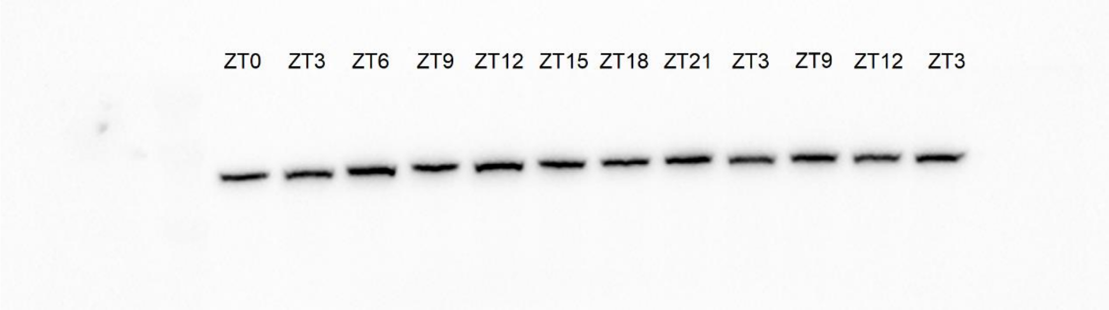


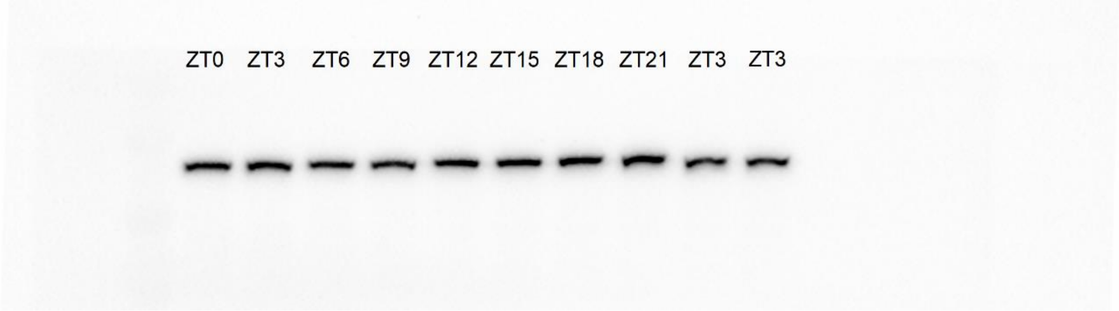


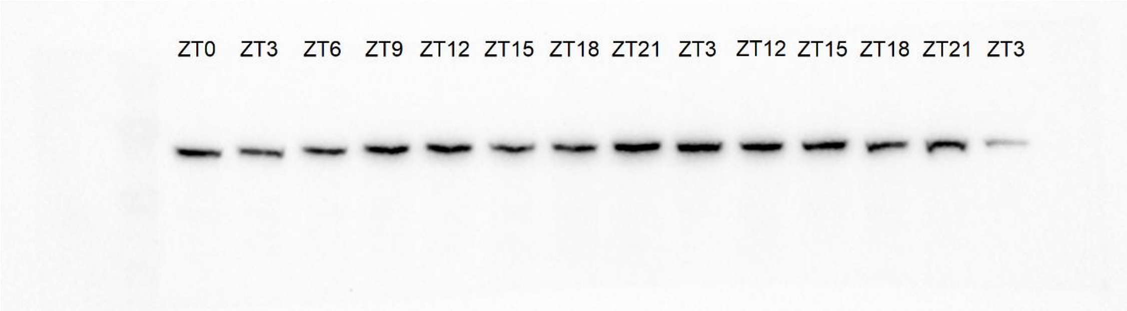

**Figure 4B, D, F, H:** Vinculin Gel 1-4 (Females)

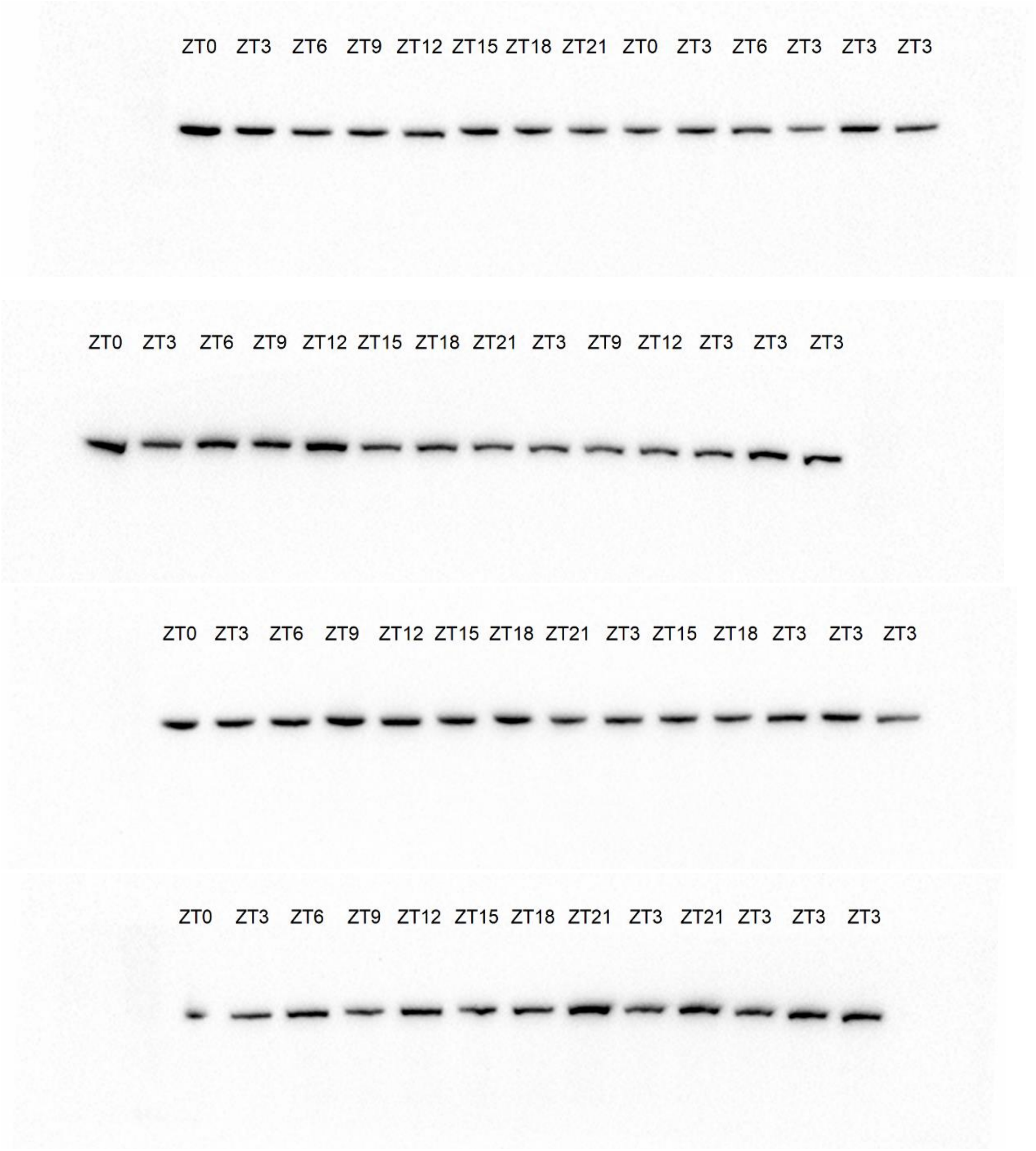
